# Keap1 recognizes EIAV early accessory protein Rev to promote antiviral defense

**DOI:** 10.1101/2021.09.29.462336

**Authors:** Yan Wang, Guanqin Ma, Xue-Feng Wang, Lei Na, Xing Guo, Jiaqi Zhang, Cong Liu, Cheng Du, Yuezhi Lin, Xiaojun Wang

## Abstract

The Nrf2/Keap1 axis plays a complex role in viral susceptibility, virus-associated inflammation and immune regulation. However, whether or how the Nrf2/Keap1 axis is involved in the interactions between equine lentiviruses and their hosts remains unclear. Here, we demonstrate that the Nrf2/Keap1 axis was activated during EIAV infection. Mechanistically, EIAV-Rev competitively binds to Keap1 and releases Nrf2 from Keap1-mediated repression, leading to the accumulation of Nrf2 in the nucleus and promoting Nrf2 responsive genes transcription. Subsequently, we demonstrated that the Nrf2/Keap1 axis represses EIAV replication via two independent molecular mechanisms: directly increasing antioxidant enzymes to promote effective cellular resistance against EIAV infection, and repression of Rev-mediated RNA transport through direct interaction between Keap1 and Rev. Together, these data suggest that activation of the Nrf2/Keap1 axis mediates a passive defensive response to combat EIAV infection. The Nrf2/Keap1 axis could be a potential target for developing the strategies for combating EIAV infection.

**Author summary:** Here we report that the Nrf2/Keap1 axis acts as an antiviral effector on EIAV replication through two-way regulation of Rev. On the one hand, EIAV-Rev activates the Nrf2/Keap1 axis by competitive interaction with Keap1, which promotes the transcription of cytoprotective genes implicated in the antiviral response. On the other hand, Keap1 restricts Rev/RRE-dependent RNA transport, leading to inhibition of viral protein synthesis and reduction in viral replication. This study highlights the importance of the Nrf2/Keap1 axis in controlling EIAV replication, and identifies this pathway as a potential new entry-point for treating lentivirus infection. Importantly, this report provides new insights into the regulation of viral replication by the Nrf2/Keap1 axis.

## Introduction

Equine infectious anemia virus (EIAV), an equine lentivirus, causes persistent infection characterized by recurring febrile episodes associated with EIA clinical signs [1, 2]. Unlike hosts with infections caused by other lentiviruses, most horses infected with EIAV become lifelong inapparent EIAV carriers by eliciting immune control over virus replication [3-5]. Additionally, an attenuated EIAV vaccine was developed through long-term passaging in equine macrophages *in vitro* and has successfully controlled the EIA epidemic in China, indicating that protective immunity can be induced after virus infection [5-7]. EIAV has therefore been widely accepted as a system for identifying potential immune control over lentiviruses [1, 8, 9]. Previous studies have shown that the initial innate immune response blocks EIAV replication and the virus has some weapons to counteract innate immunity restriction [2, 10-13]. However, the factors that determine whether infection causes death of the host or whether the host progresses to an inapparent carrier remain unknown [1, 2, 9]. Elucidation of the innate cellular defenses during EIAV infection could facilitate comprehension of the interplay between immune control and EIAV infection.

Oxidative stress is commonly induced by viral invasion, which plays a critical role in the pathogenesis of various viruses [14-17]. The activation of the antiviral and inflammatory signaling pathways has also been linked with the production of ROS [15, 17-19]. Simultaneously, the nuclear factor erythroid 2-related factor 2 (Nrf2) antioxidant pathway, which is induced downstream by ROS can be activated to maintain the cellular redox homeostasis by regulating antioxidant genes and phase II detoxification enzymes [20-22]. Moreover, several studies have demonstrated that the balance of redox homeostasis contributes to viral pathogenesis through multiple mechanisms [23-25]. The interaction of Nrf2 with its cellular inhibitor, Keap1, comprises a conserved and important intracellular antioxidant defense system [26, 27]. Under normal physiological conditions, Nrf2 is sequestered in the cytoplasm by its regulatory protein, Keap1. However, under oxidative stress, the interaction between Nrf2 and Keap1 is disrupted and the activated Nrf2 is sequentially translocated to the nucleus, where it binds the antioxidant response elements (AREs) and inducing expression of several antioxidant genes [27, 28]. As well as being involved in the anti-oxidative response, Nrf2 is also an important immune-modulator, and interferes with pro-inflammatory gene as well as innate immune signals such as TLRs, NF-ҡB, and metastasis [21, 23, 25, 28]. A growing body of evidence from studies of infection from various different viruses links the Nrf2 signal to virus replication [16, 29-32].

Recently, it has been reported that ROS and inflammatory cytokines (IL-1β and TNF-a) were generated following EIAV infection [33, 34]. Moreover, oxidative stress-induced damage and alterations in redox status have been associated with increasing disease severity in EIAV-infected horses [4]. Importantly, a transcriptomics screen as part of our previous study found that the Nrf2/Keap1 axis was activated during EIAV infection. These observations indicate a possible role for redox homeostasis in combating EIAV infection. However, whether or how EIAV infection activates the Nrf2 pathway, and the potential role of this pathway in antagonizing EIAV replication remain unclear. In the current study, we found that EIAV infection activates the Nrf2/Keap1 defense system through the EIAV accessory protein Rev. Mechanistically, Rev binds directly to the Kelch domain of Keap1, and suppresses its function, which is to inhibit Nrf2. This liberates Nrf2 and triggers the Nrf2 pathway. Moreover, the activation of Nrf2/Keap1 axis reinforces EIAV replication. We further demonstrate that the Nrf2/Keap1 axis is able to inhibit EIAV replication via two independent mechanisms: (i) enhancing antioxidant genes and (ii) the repression of Rev-mediated RNA transport, in order to reduce viral protein synthesis. In general, EIAV Rev-triggered activation of Nrf2/Keap1 axis following EIAV infection contributes to the host defenses against infection.

## Results

### EIAV Infection induces the antioxidant response and up-regulates total Nrf2 and phospho-Nrf2

To investigate the cellular response to EIAV infection, we performed a transcriptome analysis of EIAV infected equine macrophages (eMDMs). Based on the obtained transcriptome data, the observed changes in gene expression following EIAV infection were first represented using a waterfall plot, as shown in Figure 1A. More than 400 genes were either up- or down-regulated within 6 h of viral infection comparing with sham control, which indicated that most changes in gene expression occur at the early stages of infection. The subsequent analysis of intracellular signaling pathways showed that multiple canonical pathways were regulated coordinately by EIAV infection. In particular, the antiviral pathways and the inflammatory pathways were highly enriched following EIAV infection (Fig 1B). Moreover, 12 h following viral infection of cells, an enrichment in the networks associated with the oxidant stress relative response was observed (Fig 1B). Importantly, Nrf2 responsive genes including NQO1, OAS1 and HMOX1 were also up-regulated (Fig 1B and 1C), indicating the involvement of the Nrf2 pathway in the EIAV infection-mediated intracellular response. To further verify this observation, we analyzed the levels of Nrf2 in the EIAV-infected macrophages separately at both the RNA and protein levels, using real-time PCR and western blotting. The results showed that EIAV infection did indeed up-regulate levels of Nrf2 protein in a dose-dependent manner (Fig 1E and 1F), but not levels of mRNA (Fig 1G). In addition, we found that levels of total Nrf2 (tNrf2) and phosphorylated Nrf2 (pNrf2) in 293T cells transfected with an EIAV infectious clone were also elevated in a dose-dependent manner (Fig 1H).

**Fig 1.**
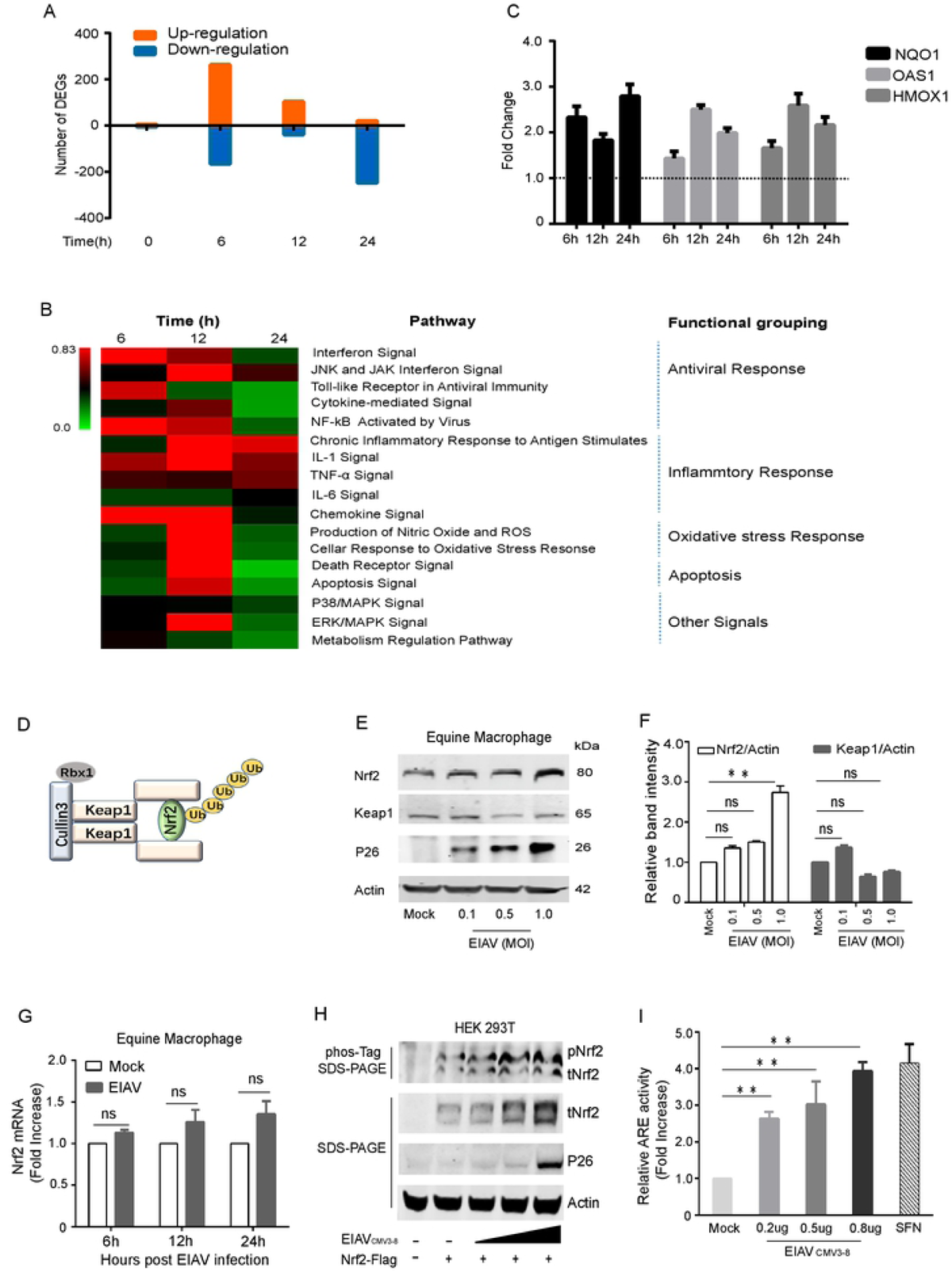
The Nrf2-Keap1 axis is activated by EIAV infection. **(A)** Waterfall plot representing the total number of up- and down-regulated genes at each time point following transcriptome analysis (DEGs, differentially expressed genes; selected based on fold change>1.5, P value<0.05). **(B)** Heat map showing statistically significant canonical pathways commonly regulated at 6 h, 12 h and 24 h post EIAV infection, compared to the control. Heat map colors represent the ratio of regulated genes after EIAV infection (red and green correspond to over- and under-regulated genes, respectively). **(C)** Real-time PCR analysis of NQO1, OAS1 and HMOX1 mRNA in equine macrophages infected with EIAV at the indicated times (6h, 2h and 24h) post infection. **(D)** Schematic representation of Nrf2-Keap1 interaction. **(E)** Quantification of Nrf2 and Keap1 expression in equine macrophages infected with EIAV at varied infection dose. **(F)** Densitometric analyses of pNrf2 and tNrf2 band intensity shown after normalization to actin. **(G)** Nrf2 mRNA was quantified using real-time PCR as above. **(H)** The effects of EIAV infection on the phosphorylation of Nrf2 were analyzed using a Phos-tag assay. Immunoblots for total Nrf2, p26 and actin were performed on normal SDS-PAGE gels as previously described. **(I)** The activation of Nrf2/Keap1 axis triggered by EIAV infection was analyzed using the ARE reporter gene assay. 293T cells were co-transfected together with the ARE luciferase reporter plasmid, as well as pcDNA3.1 (empty vector) or increasing concentrations of EIAV_CMV3-8_. Twenty-four hours later, cells were lysed and firefly luciferase activities was assayed. Data are representative of two (A-B) or three (C-I) independent experiments.

To further assess the impact of EIAV infection on the Nrf2 pathway, an ARE promotor-based luciferase reporter system was developed for the genes downstream of Nrf2. An EIAV infectious clone and the ARE-promoter reporter system were co-transfected into 293T cells. As expected, we observed more than 4-fold elevation in ARE reporter activation by EIAV clone compared to the mock-transfected control (Fig 1I). Collectively, these results demonstrated that the Nrf2 pathway was activated in EIAV-infected host cells.

### EIAV-Rev activates the Nrf2/Keap1 axis

To determine which, if any, EIAV-coded proteins are crucial for Nrf2 activation, various different amounts of expression plasmids carrying EIAV *gag, tat, s2, env*, or *rev* gene were co-transfected into 293T cells together with the luciferase reporter system. We found that expression of the EIAV-Rev protein, but not the other viral proteins, displayed a dose-dependent correlation with the ability to activate Nrf2/Keap1 axis as strongly as sulforaphane (SFN) (Fig 2A). This suggested an important role of EIAV-Rev in activating the Nrf2/Keap1 axis. To further verify Rev-triggered activation of the Nrf2/Keap1 axis, total Nrf2 (tNrf2) and phosphorylated Nrf2 (pNrf2) levels in nucleus and cytoplasm of transfected-293T cells with or without Rev were assessed separately. The expression of tNrf2 and pNrf2 were increased with presence of Rev in a dose-dependent manner both in the cytoplasm and nucleus (Fig 2B and 2C), indicating that Rev can stabilize Nrf2 and increase phosphorylation and nuclear translocation of Nrf2. Notably, suppression of Nrf2 by Keap1 was released by Rev, but not Gag or Env, resulting in the recovery of Nrf2 activation (Fig 2D). However, specific siRNA-mediated suppression of Keap1 expression promoted ARE activity to the same level when Nrf2 presented with or without Rev expression., indicating that Keap1 is the key factor for Rev mediated ARE activation (Fig 2E). Taken together, these lines of biochemical evidence indicate that EIAV-Rev is able to initiate and retain activation of the Nrf2/Keap1 axis with a Keap1-dependent mechanism.

**Fig 2.**
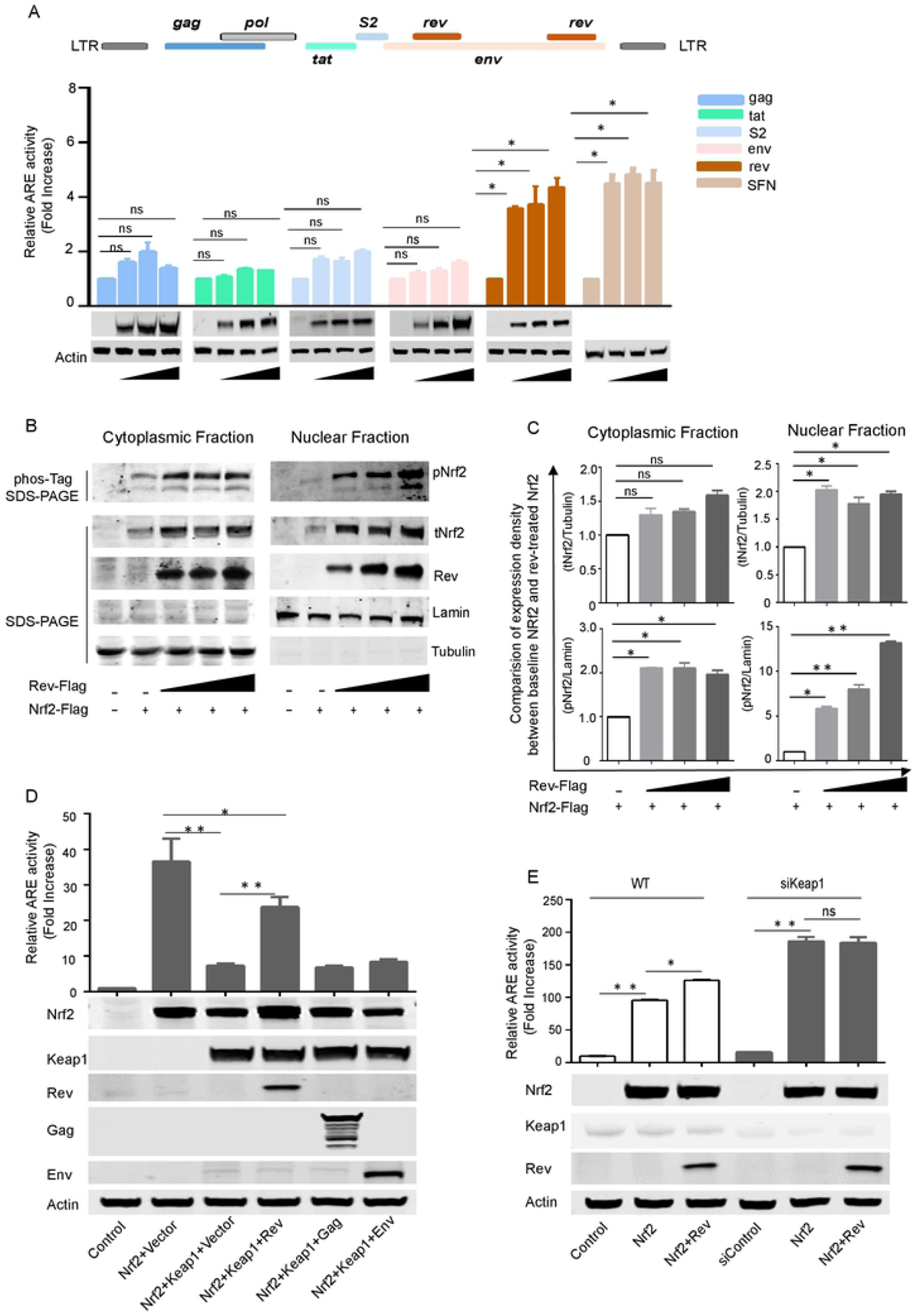
EIAV-Rev Induces Nrf2/Keap1 axis Activation. **(A)** The ability of EIAV-coded proteins inducing Nrf2/Keap1 axis activation were evaluated by ARE gene reporter. The assay protocol was the same as that shown in Figure 1I but cells were transfected with EIAV-*env*, EIAV-*gag*, EIAV-*rev*, EIAV-*S2* or EIAV-*Tat* separately. SFN was chosen as a positive control. Western blot depicting the expression of each of the transfected constructs, including the loading control, actin. **(B-C)** Rev triggered Nrf2/Keap1 axis activation. 293T cells were transfected with Flag-tagged-*rev*, and the cytoplasmic and nuclear proteins were fractionated and then immunoblotted for pNrf2 and tNrf2. Densitometric analyses of pNrf2 and tNrf2 band intensity shown after normalization to tubulin (cytoplasmic purity control) or lamin (nuclear purity control). **(D)** ARE gene reporter was assayed for the Nrf2 activation in the presence of Rev. 293T cells were transfected with the indicated plasmids. After 24 h, the luciferase activities were assessed (upper) and exogenous expression of proteins was measured using western blotting (lower). **(E)** Same assay protocol as D but 293T cells were pre-treated with siControl or siKeap1 and then transfected with the indicated plasmids. Data shown represent three independent experiments. Data are the mean values ± SDs, ns (non-significant), P > 0.05; *P < 0.05; **P < 0.01 (Student’s t test)

### EIAV-Rev interacts with Keap1 through its Kelch domain

To investigate how Rev induces the activation of the Nrf2/Keap1 axis, we first determined whether there was a physical interaction between Keap1, Nrf2 and Rev using a proximity ligation assay (PLA) and co-immunoprecipitation (Co-IP) assays. The results showed that Rev has a direct interaction with Keap1 (HA-tagged) but not Nrf2 (HA-tagged) (Fig 3A-C). Next, the crucial domain on Keap1 necessary for the interaction with Rev was also identified using Co-IP assay, with co-transfection of Rev together with various Keap1 mutants, including N-terminal region (NTR), Bric-a-Bric domain (BTB), intervening region (IVR), Kelch domain, and C-terminal domain (CTR) deletions (Fig 3D). Co-IP results showed that Keap1 mutants lacking NTR, BTB, IVR or CTR were precipitated with Rev similarly to the full length Keap1 (Fig 3E). However, the Keap1 mutant without Kelch domain completely lost the ability to bind Rev (Fig 3E), suggesting that the Kelch domain is required to interact with EIAV-Rev. Similarly, this Keap1 mutant (Kelch domain deletion) did not co-precipitate with Nrf2, consistent with previous studies [30], indicating that Nrf2 and Rev interact with Keap1 through a similar region (Fig 3F).

**Fig 3.**
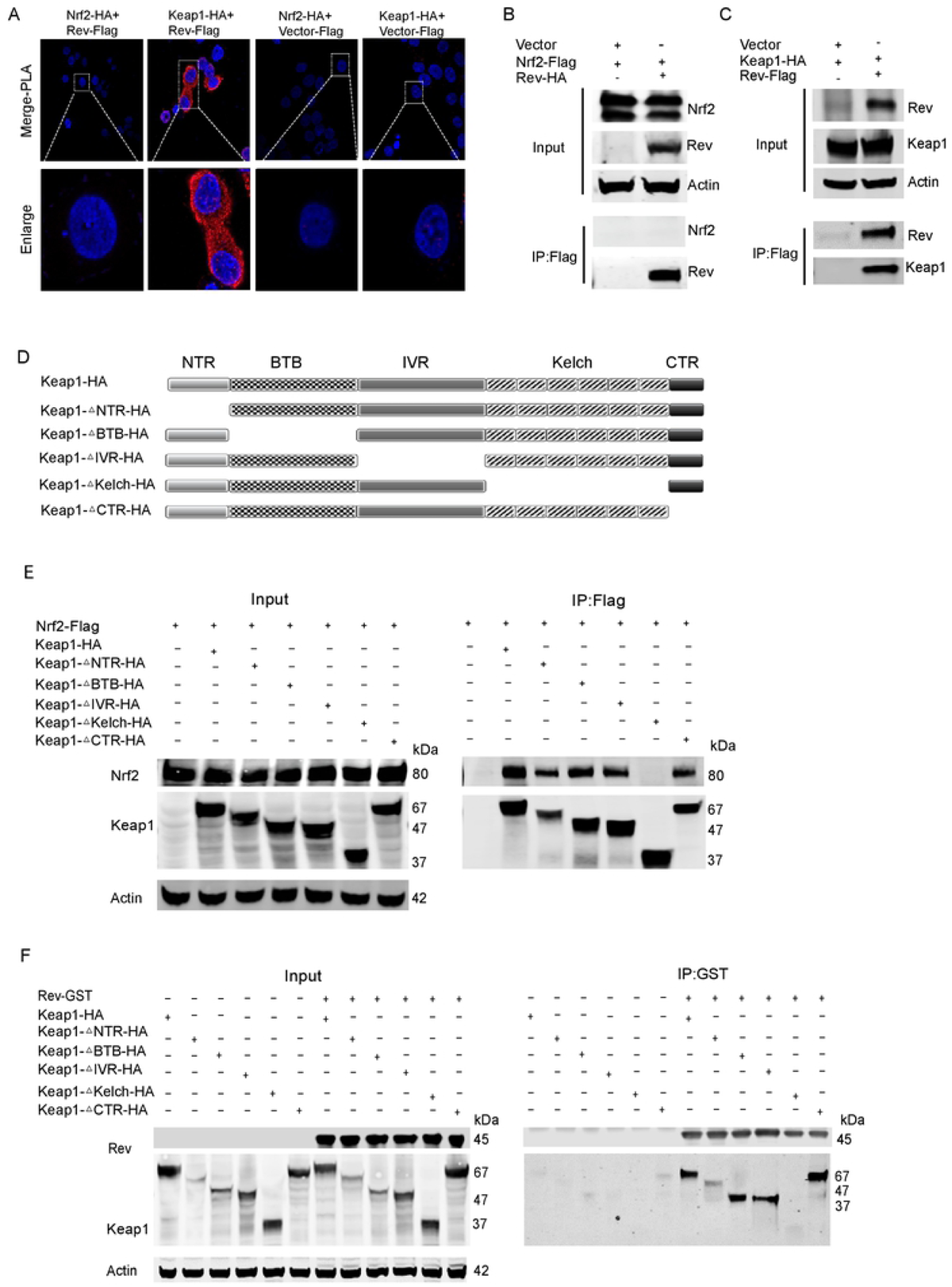
EIAV-Rev interacts with Keap1 but not Nrf2. **(A)** PLA assay was used to analyze the interactions between Rev and Keap1 or Nrf2. The red spots represent interacting complexes of the examined proteins. The nuclei were stained with DAPI (blue). Cells co-transfected with Keap1 (Nrf2) and vector were used as negative controls. **(B)** and **(C)** Reciprocal immunoprecipitations of Rev and Keap1 or Nrf2 were performed on 293T cells co-transfected with plasmids for Nrf2-Flag and *rev*-HA or with Keap1-HA and Flag-*rev*. 24 hours post-transfection, cells were collected and subjected to pull down with Flag beads. **(D)** Schematic diagram of Keap1 domain structure and Keap1 deletion mutants used in (E and F). All Keap1 deletion mutants were HA tagged at the amino terminal end. **(E)** The interaction of Keap1 mutants and Nrf2 were screened by Co-IP with the selected antibodies. **(F)** as **(E)** with additional of GST-tagged-*rev* instead of Nrf2-Flag.

Previous work has suggested that the “DLG” and “ETGE” motifs located in the Nrf2 Neh2 region were required for the binding of Nrf2 and Keap1. By screening Rev amino acid sequences, we also found a “GE” motif (S1 Fig A). We therefore sought to discover whether Keap1 interacts with Rev in a similar manner as with Nrf2. To this end, Rev mutants were developed as shown in Figure 4A defective in either the “DYG” or “RWGE” motifs singly, or defective in both. We then co-expressed Flag-tagged Rev mutants together with HA-tagged Keap1 in 293T cells and assessed the binding between these two proteins using Flag antibody-based Co-IP. Interestingly, these Rev mutants all still possessed the capacity to interact with Keap1 (S1 Fig B). These data suggested that there might be another specific sequence on Rev, with the ability to bind to Keap1. To gain further insights into the domain (s) of Rev responsible for the interaction with Keap1, we designed three segmental Rev mutants with different area deletions (S1 Fig C and D). Co-IP assays showed that an N-terminal (1-56aa) deletion on Rev led to failure of the interaction between Rev and Keap1. Moreover, this N-terminal deletion also resulted in Rev losing the ability to activate the Nrf2/Keap1 axis (S1 Fig E). Collectively, these data indicated that 1-56aa at the Rev N-terminal is the key region both for binding Keap1 and activating the Nrf2/Keap1 axis.

### Competitive binding of Nrf2 or Rev to Keap1 in 293T cells

If Nrf2 and Rev both interact with the Keap1 Kelch domain, it is possible that binding of Nrf2 and Rev to Keap1 is competitive. To address this question, we performed a biolayer interferometry (ForteBio) assay to test affinity between Keap1, Nrf2 and Rev. The results showed binding between Keap1 and Rev and between Keap1 and Nrf2, regardless of which protein was tethered to the sensor. However, no interaction between Nrf2 and Rev was observed (data not shown). This biophysical binding result was consistent with the Co-IP results. The binding curves of Keap1-Nrf2 and Keap1-Rev were found to be very similar and with comparable biophysical affinity (KDM Keap1-Nrf2 = 3.995E^-07^ vs KDM Keap1-Rev = 4.9375E^-07^) (Fig 4A and 4B). The binding between Keap1 and Nrf2 in either the presence or absence of Rev was then examined, to determine the competitive effect of Rev to Nrf2 when binding to Keap1. The results showed that it was difficult for Keap1 to interact with Nrf2 (no increasing binding signal was observed for the blue line) when Keap1 and Rev were pre-combined (saturated state), while a significant binding signal was observed in the control group (Keap1-buffer-Nrf2) without pre-binding of Rev and Keap1 (Fig 4C and 4D). Similar binding dynamics were observed when the binding of Keap1 and Rev with or without pre-combination of Keap1 and Nrf2 was tested. Taken together, these results confirmed that Rev and Nrf2 bind competitively to Keap1 *in vitro*.

**Fig 4.**
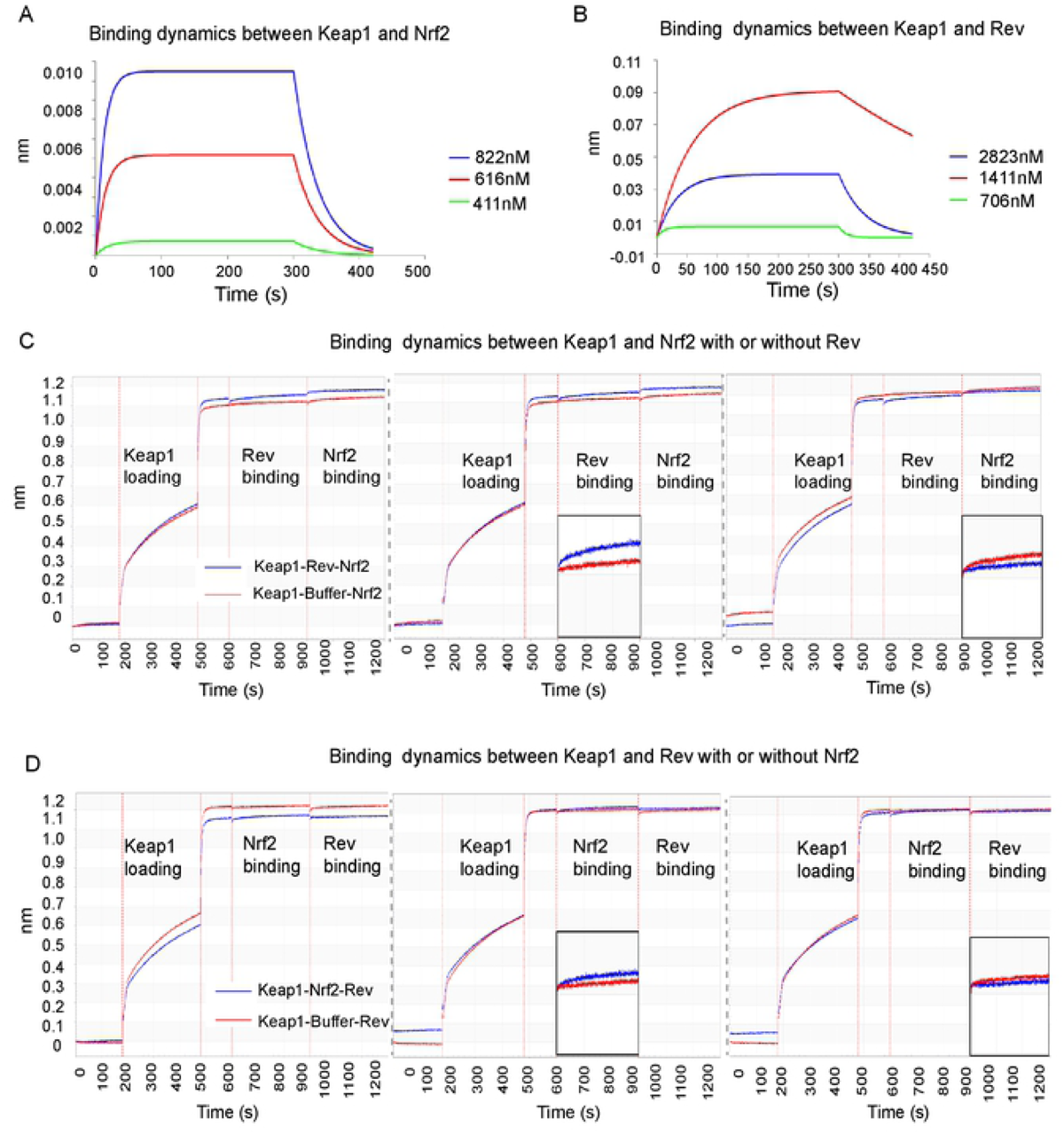
Rev-competitive inhibition antagonizes Nrf2 binding to Keap1. **(A-B)** Biolayer Interferometry graphs showing association and dissociation steps using different concentrations of Nrf2 (A) or Rev (B) to Keap1 immobilized on the anti-streptavidin biosensors. **(C-D)** BLI profiles showing the effect of Rev on the affinity of the Keap1-Nrf2 interaction. Rev incubated with Keap1 pre-coated with Nrf2 on the biosensors (C). Nrf2 (blue line) incubated with Keap1 pre-coated with Rev (D). The data are expressed as means and SD for at least three independent replicates.

### EIAV-Rev prevents Keap1-mediated Nrf2 degradation and enhances Nrf2 nuclei translocation

We have shown that Rev binds to Keap1 and impairs the interaction between Keap1 and Nrf2. Given that Keap1 is known to mediate Nrf2 degradation via ubiquitination [35], we next tested whether Rev affects the status of Keap1-mediated Nrf2 degradation. First, Keap1 and Nrf2 were co-transfected into 293T cells with or without Rev. As expected, Rev was able to reverse Keap1-mediated degradation of Nrf2 and restore the levels of Nrf2 in the cytoplasm (Fig 5A, compare lanes 2 and 3). Moreover, overexpression of Rev resulted in a significant decrease in Keap1-mediated Nrf2 ubiquitination (Fig 5B). As EIAV infection did not impact Keap1 protein levels in host cells (Fig 1D and 1E), all these data indicated that the increased Nrf2 was as a result of the reduction of Keap1-mediated Nrf2 degradation regulated by Rev. To provide further evidence of the ability of EIAV-Rev to release Nrf2 from its negative regulator Keap1, we investigated the distribution of Nrf2 with or without Rev co-expression with an immunofluorescence assay. When Nrf2 was ectopically expressed alone, it showed a nuclear distribution, whereas it exhibited a homogenous cytoplasmic distribution if co-expressed with Keap1 (Fig 5D). Importantly, in cells co-expressing Nrf2 and Keap1, the addition of Rev resulted in translocation of Nrf2 from the cytoplasm to the nucleus (Fig 5E). These results further suggest that the interaction between Rev and Keap1 inhibits Keap1-mediated degradation of Nrf2, leading to the translocation of Nrf2 from the cytoplasm to the nucleus.

**Fig 5.**
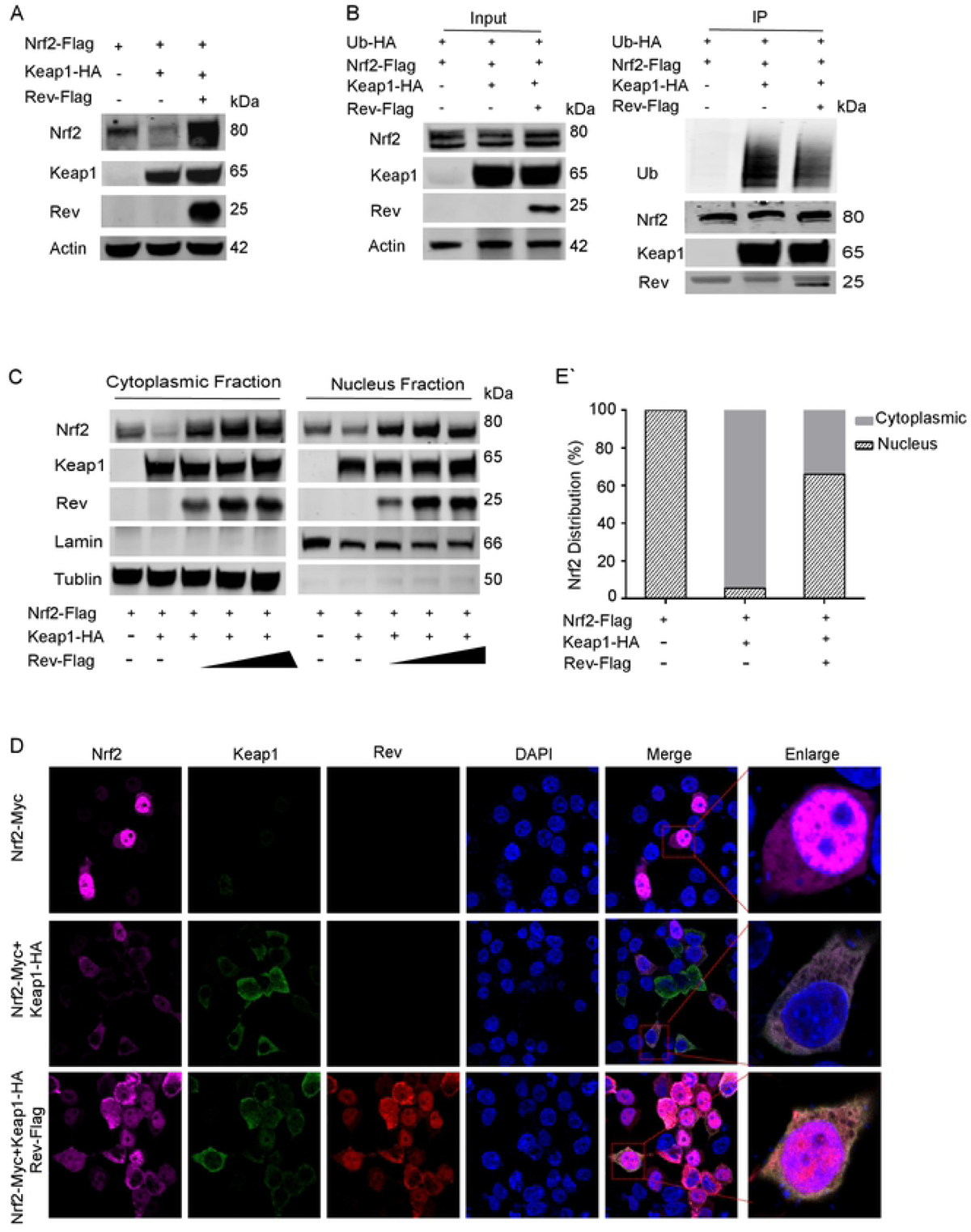
EAIV-Rev reduces Keap1-mediated Nrf2 ubiquitination and facilitates Nrf2 Translocation into the Nuclei. **(A)** Rev prevents Keap1-mediated Nrf2 degradation. Keap1 and *rev* were expressed in 293T cells alone or in combination as indicated. Cells were analyzed by western blot at 24 h p.t. **(B)** Rev reduces Keap1-mediated Nrf2 ubiquitination. 293T cells were transfected with the indicated plasmids. At 24 h post-transfection, whole-cell lysates were immunoprecipitated by Flag beads and then ubiquitination was analyzed by blotting with HA-K48 ubiquitin antibody. **(C)** Protocol as in (A), with the addition of a step separating the cytoplasmic and nuclear proteins. Western blotting was performed as in Figure 3B. **(D)** The distribution of Nrf2 was visualized with or without Rev using confocal imaging. The plasmids encoding Nrf2 (purple) and Keap1 (green) were expressed alone or in combination with *rev* (red) in 293T cells. Twenty-four hours later, cells were fixed using acetone/methanol and subjected to immunofluorescence analysis using the indicated antibodies. Nuclei were visualized by staining with DAPI. Quantifications are given in **(E)**. In each transfection experiment, at least 100 cells were scored, and the gray and stripe bars represent cytosolic and nuclear localization, respectively. Each of these experiments was repeated at least twice, and consistent results were obtained.

### Nrf2 inhibits EIAV virion production in equine macrophages and 293T cells

So far, our data supported the idea that EIAV-Rev promotes Nrf2 function via competitively binding to Keap1. However, the role of Rev-mediated Nrf2 activation in EIAV production and replication remains unclear. To investigate this, we first knocked out the Nrf2 gene in 293T cells (Nrf2_ko_) and used an EIAV infectious molecular clone to transduce Nrf2_ko_ 293T cells. We found that the production of virions was significantly increased in cell lysate and supernatant of Nrf2_ko_ cells compared to the control 293T cells. Moreover, the increased virion production in Nrf2_ko_ cells was repressed when Nrf2 protein was re-constituted (S2 Fig A and B). Furthermore, ectopic expression of Nrf2 in 293T cells repressed viral production in a dose-dependent manner (S2 Fig C and D). Over expression of Keap1 could down regulate Nrf2 expression and enhance viral production (S2 Fig E and F). These results indicated that Nrf2 play important role in anti-EIAV replication. To further confirm this observation, we used equine macrophages, the EIAV natural host cells, to verify the interaction between EIAV and the Nrf2 pathway. We utilized Nrf2-specific siRNA to effectively decrease Nrf2 expression in equine macrophages and infected these eMDMs with EIAV. As shown in Fig 6A, we did see EIAV-P26 protein level was up-regulated by ∼4 folds in Nrf2 KD eMDMs compared to Si-control RNA-treated eMDMs. We also observed the enhanced virus P26 protein production in cultural supernatant (Fig 6B), which is consistent with up-regulated intracellular EIAV Gag expression. As previous studies have shown, the compound SFN can activate the Nrf2 pathway. We pre-treated eMDMs with 20 μM SFN for 6 h to activate the Nrf2 pathway (as assessed via increases in HO-1 protein), and then infected them with EIAV. Western blot assays showed detected decreased virion production in either cell lysates or supernatants (Fig 6C and 6D). After we confirmed the ability of the Rev-triggered Nrf2 pathway to regulate EIAV infection, we next used a VSV-G pseudotyped *env* (‒) EIAV luciferase reporter virus system to investigate whether Rev was able to serve as Nrf2-activator to suppress virus infection. The results showed that EIAV infection was obviously repressed with over-expression of Rev (Fig 6E). Most interestingly, the HIV-1 luciferase reporter system showed similar response (Fig 6F). Together with the result from figure 4 and figure5, these data demonstrated that Keap1 could recognize and bind Rev to release Keap1, and further activate antiviral response to block EIAV and HIV-1 replication.

**Fig 6.**
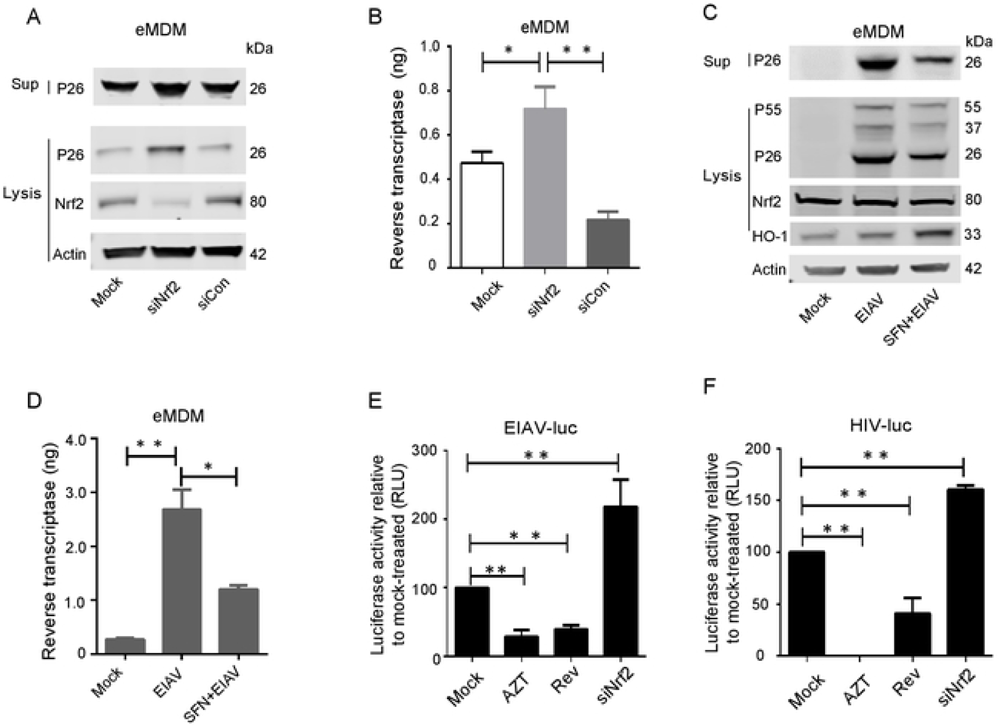
Nrf2 blocks EIAV replication in equine macrophages. **(A-B)** EIAV replication was determined in eMDM cells treated with Nrf2-specific siRNA or control RNA. Viral protein expression in supernatant was calculated as described as B. **(C-D)** eMDMs were treated with media supplemented with 20 μM SFN. Six-hours after treatment, cells were either mock infected or infected with EIAV at 1×10^5^ TCID50. 24 hours post infection, cell extracts were analyzed using western blotting and reverse transcriptase activity assay. **(E-F)** 293T cells were transfected with *rev* or Nrf2-specific siRNA. After treatment for 12h, cells were inoculated with VSV-G-pseudotyped EIAV encoding firefly luciferase (K) and VSV-G-pseudotyped HIV-1 (L). Twenty-four hours later, luciferase activity was measured using photon emission. Zidovudine (AZT), a specific reverse transcription inhibitor, served as a positive control for viral inhibition. The data are expressed as means and SD for at least three independent replicates. *P <0.05, * *P<0.01.

### Keap1 limited Rev-mediated viral mRNA export correlated with low virus production

Having confirmed the reduced viral production resulting from Nrf2/Keap1 activation induced by Rev, we next ask whether the binding of Rev by Keap1 could also directly inhibit viral protein expression, as EIAV replication is known to be absolutely dependent upon Rev-mediated transport of viral RNA [36, 37]. We first investigated the sub-cellular localization of Rev and Keap1. Our results suggested that Rev was located predominantly in the nucleus in the absence of Keap1, while in the presence of Keap1, Rev was trans-located mostly in the cytoplasm and co-localized with Keap1 (Fig 7A and 7B). Subsequently, EIAV pseudotyped virus was used to infect siKeap1- or scrambled siRNA-treated cells. The nuclear and cytoplasmic RNA fraction were then isolated and purified, and the cytoplasmic and nuclear distributions of EIAV-*gag* mRNA were analyzed using quantitative PCR at 12 h and 24 h post infection. Our results showed that most of the *gag* mRNA had accumulated in the cytoplasmic when Keap1 endogenous expression was silenced (Fig 7C). These data indicated that the efficiency of Rev-mediated RNA transport between the cytoplasm and nucleus was greatly enhanced in siKeap1 cells (Fig 7C). As a result of the increased transport of the primary *gag* transcript, viral production was higher in siKeap1 cells, compared with control cells under transfection and infection conditions (Fig 7D and 7E). These data indicated that Keap1 may limit Rev-mediated RNA transport through direct interaction with Rev. To investigate this phenomenon further, we compared the effects of Keap1 on RNA transported by different RNA export systems. We found that RNA transport was promoted in the Rev-dependent Rev/RRE export system (Rev-dependent) in the absence of Keap1. In contrast, there was no effect in the Rev-independent 4xCTE export system (Fig 7F and 7G). We therefore surmised that binding with Keap1 could lead to the accumulation of Rev in the cytoplasm and subsequently impair Rev/RRE dependent RNA transport, inhibiting *gag* mRNA production in trans and leading to reduced viral production. Taken together, these data indicated the dual roles of Keap1 in the antiviral response: 1) the sensing viral Rev protein to activate Naf2-AREs pathway; 2) the direct binding and block of Rev to inhibit viral RNA transportation.

**Fig 7.**
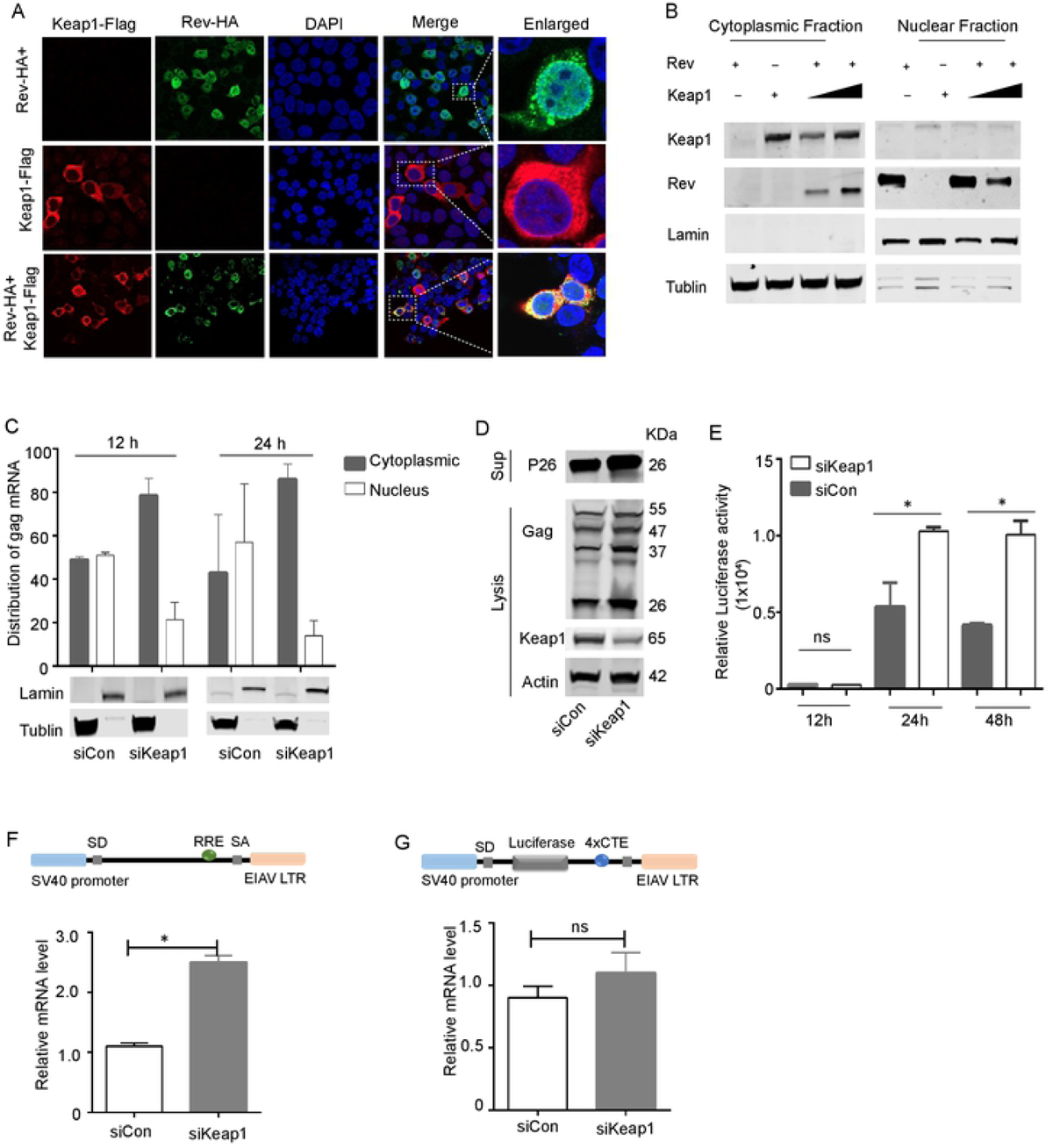
Keap1 hijacks *rev* in the cytoplasm and limits *rev*-mediated RNA transport. **(A)** Distribution of Rev was screened with or without Keap1 by confocal imaging. Scale bars 10µm. Images are representative of 3 independent experiments. **(B)** 293T cells were co-transfected with HA-tagged-*rev* and Keap1 at dose. The cytoplasmic and nuclear proteins were fractionated as in Figure 2B and then immunoblotted for Keap1 and Rev. **(C)** Distribution of cytoplasmic and nuclear unspliced *gag* RNAs in siKeap1 and control cells was analyzed using real-time PCR with primers specific for *gag* mRNA. Lamin and tubulin were used as nuclear and cytoplasmic controls, respectively. **(D)** 293T cells were treated with siRNA targeting Keap1 (siKeap1) or scrambled siRNA (siControl) and then transfected with EIAV_CMV3-8_. Cells were lysed and samples were analyzed for expression of Gag, Keap1, and Actin using western blotting. **(E)** EIAV pseudovirions were generated separately in siKeap1 or siControl cells and then used to infect 293T cells. The luciferase activity in the supernatant was assayed at the indicated timepoints (12h, 24h and 48h). (**F)** Rev/RRE RNA export reporter plasmids were transfected into siKeap1 and control cells and viral mRNA synthesis was calculated using real-time RT PCR. **(G)** The protocol is as (F), but with the Rev-independent RNA export reporter plasmid (4XCTE). The graph represents three independent experiments; error bars represents SEM.

## Discussion

Activation of the Nrf2/Keap1 axis through various mechanisms to regulate oxidative stress has been described from the infection of several viruses, including hepatitis C virus (HCV), HBV, influenza and HIV [31, 38-40]. However, the mechanisms that underline Nrf2/Keap1 axis activation, especially the roles of Keap1 in antiviral defense, have not been fully elucidated. Here, we revealed a novel antiviral mechanism by which the Nrf2/Keap1 axis was activated following EIAV infection and proposed a working model of Keap1 (Fig 8). Briefly, Keap1, a specific repressor of Nrf2, can directly bind to EIAV-Rev upon EIAV infection as a sensor, and disrupt the binding of Keap1 and Nrf2, leading to releasing and nuclear localization of Nrf2 and triggering the antioxidant response. At same time, as an effector, Keap1 binds and retains Rev in cytoplasm, disrupts the Rev’s function in transportation of viral RNA, and further represses EIAV production in host cells.

**Fig 8.**
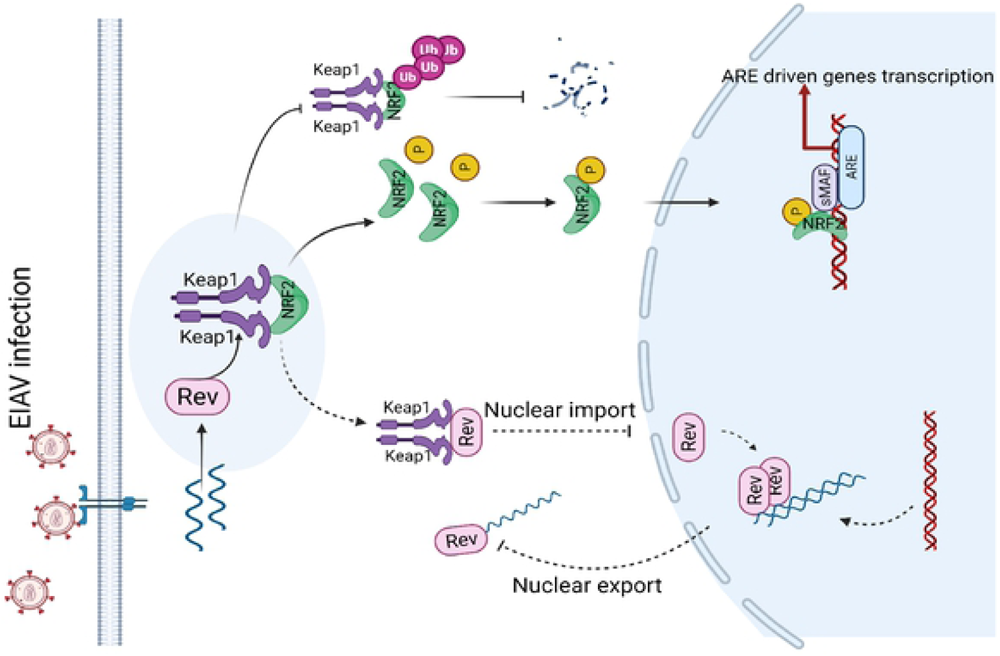
A proposed model for the cellular Nrf2/Keap1 axis manipulation of EIAV replication. Nrf2/Keap1 plays a crucial role in the management of oxidative stress. Under normal conditions, Keap1 interacts with Nrf2 in cytoplasm, targeting it for proteasomal degradation to maintain redox homeostasis. Under EIAV infection, EIAV-Rev interacts with Keap1, disrupting the interaction of Keap1-Nrf2, leading to the enhancement of pNrf2 and nuclear translocation which results in the activation of Nrf2. The activation of Nrf2 signal in turn inhibits EIAV replication via increasing expression of several antioxidant enzymes and also by limiting *rev*-mediated RNA transport.

Many viral infections can cause host cellular oxidative stress and further induce Nrf2 activation. Interestingly, some viruses such as Kaposi’s Sarcoma-Associated Herpesvirus (KSHV) infection benefit from Nrf2 activation [41-43]. However, Dengue virus and Respiratory syncytial virus (RSV) infection increases reactive oxygen species (ROS) and induce degradation of Nrf2 by different mechanisms [44-46]. As the key molecule that regulates the Nrf2-ARE pathway, Keap1 binds to Nrf2 to maintain a cellular antioxidant defense homeostasis. Few is known on the role of Keap1 in regulation of viral infection except that Marburg virus (MARV) VP24 targets Keap1 to drive activation of the Nrf2, which likely contribute to viral infection [30, 47]. Previous study showed that HIV-1 Tat protein can activate Nrf2 pathway which in turn suppress Tat-induced HIV-1 LTR transactivation [48]. Here we report novel roles of Keap1 in the defense of equine lentivirus EIAV infection by directly targeting viral accessory protein Rev. The pattern that the direct binding of Rev by Keap1 inhibits viral RNA transportation is unique and has not been seen in other viruses. This evidence proposed that the cellular Keap1/Nrf2 pathway was able to utilize virus-coded protein as a trigger to exert anti-viral defenses.

The Keap1-Nrf2 interaction is required for negative regulation of the anti-oxidant pathway [49]. We find that EIAV Rev binds to Keap1’s Kelch domain, which is the same domain that used to bind Nrf2. With similar binding affinity and dynamics of Keap1 to Rev and Nrf2, the recognizing and interacting with Rev by Keap1 results in releasing of Nrf2 from Keap1, subsequently Nrf2 is accumulated in the nucleus and increases Nrf2-dependent ARE activity. Two types of Nrf2/Keap1 activation, the “canonical” and “non-canonical” mechanisms, have been documented [49, 50]. Oxidative or electrophilic compounds cause electrophilic-modification of one or more cysteine in Keap1, which results in a conformational change that reduces the ability of Keap1 to bind with Nrf2 [49, 50]. This is known as the “canonical” activation pattern [49]. Here, we found that EIAV Rev could disrupt the Keap1-Nrf2 interaction through direct interaction with Keap1. Hence, the Nrf2 activation induced by EIAV Rev belongs to the “non-canonical” pattern, through disruption of the Keap1-Nrf2 complex via interaction with Keap1 or Nrf2 [49, 50]. Several cellular factors, including P62, BRC interaction A1 and DPP3, are known to be able to reduce or abolish the binding of Keap1 and Nrf2 in this way [49]. However, except for Marburg virus VP24 [47], it has only rarely been reported that virus-coded proteins involved in the regulation of the Nrf2/Keap1 axis regulation act through direct interaction.

Keap1 interactors including Nrf2, p62, WTX, and prothymosin alpha interact with Keap1 arginine residues mainly through electrostatic interactions [51-53]. Keap1 interactors also displayed acidic residues along with a “GE motif” [54, 55]. It has been confirmed that Keap1 binding to Nrf2 occurs via two recognition sites (the “ETGE” and “DLG” motifs), forming a “hinge-latch” like structure [55]. Keap1-mediated repression of Nrf2 is dependent on this structure [54]. Given that a “GE motif”-like sequence was found at the Rev C-terminal, we speculated that the interaction of Keap1 and Rev has a similar mechanism as the Keap1-Nrf2 interaction. However, the mutation of these two motifs to alanine was not sufficient to reduce or abrogate the binding of Rev and Keap1. Importantly, the N terminal (1-56aa) of *rev* was proved to be necessary for its binding with Keap1, as well as Keap1-dependent Nrf2 activation, without exhibiting any similarity with “GE motifs”. These discoveries suggest that Rev might present other motifs for interaction with Keap1. It has been reported that “non-covalent” Nrf2 activators could be potential candidates to break up the Keap1-Nrf2 complex due to their specificity or the ability to avoid prolonged Nrf2 activation [50, 56, 57]. Because of this, some peptides derived from the “ETGE” motif of Nrf2 or some other small molecules have the ability to disrupt the Keap1-Nrf2 complex, making Nrf2-targeted therapeutics possible [22, 27, 57]. Therefore, uncovering the residues responsible for the interaction between Rev and Keap1 will also benefit exploration for novel drugs for Nrf2-targeted therapeutics.

Taken together, the findings reported here provide insights into the molecular mechanism underlying viral infection induced Nrf2/Keap1 axis activation and reveal a novel paradigm for how cellular pathways can target virus components to facilitate anti-virus defense functions. Additionally, our results identify potential domain sequences that are responsible for Keap1 binding and triggering the Nrf2/Keap1 axis to control EIAV replication, and even also for HIV-1. The current studies highlight the significance of the Nrf2/Keap1 axis in the regulation of EIAV replication and broaden our understanding of antioxidant-stress induced by virus infection.

## Materials and Methods

### Reagents and Antibodies

Sulforaphane (SFN) (LKT Laboratories, S8044) was bought from MedChem Express and diluted with dimethyl sulfoxide (DMSO). Azidothymidine (AZT) (LKT Laboratories, S8044) was bought from MedChem Express and dissolved in water. The mouse anti-actin (A1978), mouse anti-Flag (F1804), and mouse anti-HA (H9658) monoclonal antibodies; the rabbit anti-Flag (F7425), rabbit anti-HA (H6908), and anti-Mouse IgG-FITC antibodies were purchased from Sigma. Rat anti-Myc, mouse anti-Tubulin (TA503129) monoclonal antibodies, Rabbit anti-Lamin (EPR8985), Alexa Fluor 568-conjugated (ab175476) and Alexa Fluor 647-conjugated (ab190565) antibodies were purchased from Abcam (Cambridge, UK). The rabbit anti-GST (10000-0-AP) antibodies were purchased from Proteintech. DyLight™ 800-labeled goat anti-mouse (5230-0415) and DyLight™ 508 680-labeled goat anti-rabbit (5230-0403) secondary antibodies were purchased from KPL. Anti-eqNrf2, anti-eqKeap1 and anti-P26 antibodies were prepared in our laboratory. The nuclear extraction kit was purchased from Invent Biotechnologies (SC-003). Non-cytotoxic dosages were used in this study.

### Plasmids construction, transfection and Infection

The eqNrf2 and eqKeap1 cDNA used in this study were cloned from equine monocyte-derived macrophages (eMDMs) using RT-PCR with the following set of primers:

eqNrf2 sense, 5’- ATGATGGACTTGGAGGTGC -3’;

eqNrf2 anti-sense, 5’- CTTCTTCTTGACGTCTGTCTTC-3’;

eqKeap1 sense, 5’- ATGATGATGTTCGCGTCCACC-3’;

eqKeap1 anti-sense, 5’- CTTCTTGACGTCTGTCTTC-3’;

The eqNrf2 constructs were obtained by cloning PCR-generated fragments into the VR1012 vector tagged with Flag or Myc at the N-terminal. The eqKeap1 expression vector was inserted into a pcDNA3.1 expression vector with an HA tag at the C-terminal. All expression vectors used the recombinant DNA techniques used the In-Fusion HD enzyme (Clontech, Felicia, CA). A series of deletion mutants of eqkeap1 (eqKeap1-ΔNTR, eqKeap1-ΔBTB, eqKeap1-ΔIVR, eqKeap1-Δkelch, eqKeap1-ΔCTR-HA) were generated using standard oligo-directed mutagenesis techniques. Constructs of *rev* and its mutants were created using the same techniques. For transfections, the corresponding plasmids were transfected using Poly Jet™ transfection reagent (SignaGen Laboratories, SL100688) after the 293T cells had been grown to 80% confluence for 18-24 h before treatments. Cell lysates and culture supernatants were collected 24 hours after transfection. For infection, the equine monocytes were infected with EIAV at 1×10^5^ TCID50. Cell extracts were collected and analyzed according to the requirements of the experiment. Details of plasmid constructions are provided in the supplemental experimental procedures.

### Microarray analysis

Equine macrophages were infected at an MOI of 10 as described above, cells were collected at the indicated times and TRIzol reagent (Qiagen) was used for RNA extraction, according to the manufacturer’s protocol. RNA purity and quantification were evaluated using a NanoDrop 2000 spectrophotometer (Thermo Scientific, USA). RNA integrity was assessed using an Agilent 2100 Bioanalyzer (Agilent Technologies, Santa Clara, CA, USA). Libraries were then constructed using TruSeq Stranded mRNA LT Sample Prep Kit (Illumina, San Diego, CA, USA) according to the manufacturer’s instructions. Transcriptome sequencing and analysis were performed by OE Biotech Co., Ltd. (Shanghai, China). The libraries were sequenced on an Illumina HiSeq X Ten platform and 150 bp paired-end reads were generated. About 59437095 raw reads for each sample were generated. Raw data (raw reads) of fastq format were first processed using Trimmomatic [58] and the low quality reads were removed to obtain clean reads. About 58897197 clean reads for each sample were retained for subsequent analyses. The clean reads were mapped to the human genome (GRCh38) using HISAT2. FPKM of each gene was calculated using Cufflinks, and the read counts for each gene were obtained by HTSeq-count [59]. Differential expression analysis was performed using the DESeq (2012) package in R [60]. A p value < 0.05 and fold change > 2 or fold change < 0.5 was set as the threshold for significantly differential expression. Hierarchical cluster analysis of differentially expressed genes (DEGs) was performed to demonstrate the expression patterns of genes in different groups and samples. GO enrichment and KEGG pathway enrichment analysis of DEGs were performed separately using R, based on the hypergeometric distribution. The pathway enrichment and network analyses were performed based on this calculation: ratio of the number of up-regulated genes to the total number of genes that map to the canonical pathway.

### Activation of the Nrf2/Keap1 axis

For ARE reporter gene assays, 293T cells were co-transfected with the plasmids expressing ARE-inducible firefly luciferase (pARE-Luc; Beyotime) and the following plasmids expressing viral proteins: VR1012-*gag* /pcDNA3.1-*env* /pcDNA3.1-*S2* /pcDNA3.1-*Tat* /VR1012-*rev* separately, using the Poly Jet™ transfection reagent as previously described. After 24h, a dual luciferase reporter assay (Promega, E1500) was performed in triplicate and firefly luciferase values were normalized to *Renilla* luciferase values. The expression of the corresponding target proteins was detected using western blotting. Levels of endogenous mRNAs from Nrf2 and Nrf2-downstream genes (NQO1, OAS1 and HMOX1) were assessed using quantitative real-time RT-PCR with 2×SYBR Green Fast qRT-PCR Master Mix (Biotool). The value obtained for each gene was normalized to β-actin expression. The sequences of primers used in this study were as follows:

eqNrf2 sense: 5’-ATGGATTTGATTGACAT-3’;

eqNrf2 anti-sense: 5’-TCACCTGTCTTCATCTAGTT-3’;

eqNQO1 sense: 5’-TGGAAGGATGGAAGAAACG-3’;

eqNQO1 anti-sense: 5’-GGACTTGCCCAAGTGATG-3’;

eqOAS1 sense: 5’-AGGCTACCCCAATATGAATCA-3’;

eqOAS1 anti-sense: 5’-ATCCCCAAGGCCCATC-3’;

eqHMOX1 sense: 5’-CCCAGGATTTGTCAGAGGCC-3’;

eqHMOX1 anti-sense: 5’-TGGTACAGGGAGGCCATCAC-3’.

### Immunoprecipitation, immunoblot analysis, and ubiquitination assay

These experiments were performed as previously described. For immunoprecipitation, samples were prepared after transfection with the indicated plasmids, and then incubated overnight with the appropriate antibodies together with anti-HA magnetic beads (Sigma-Aldrich, A2095), anti-Flag magnetic beads (Sigma-Aldrich, A2095) or anti-GST magnetic beads (Genscript, L00327). The beads were washed three to five times with ice-cold PBS and incubated with the cell lysates overnight at 4°C.The immunoprecipitates were washed three times in 1 ml SDS lysis buffer and subjected to immunoblot analysis. For *in vitro* ubiquitination experiments, 293T cells were transfected with the indicated plasmids for 48 hr. Cells were lysed under denaturing conditions in an SDS buffer (50 mM Tris-HCl, pH 7.5, 0.5 mM EDTA, 1 mM DTT, 1% SDS) by boiling for 10 min. The lysate was subjected to immunoprecipitation using anti-Flag magnetic beads and subsequent SDS-PAGE. Ubiquitylated Nrf2 was detected with anti-HA antibody. The protein band was detected using a Licor Odyssey imaging system (USA) and quantified using ImageJ software.

### Proximity ligation assay (PLA)

A proximity ligation assay (PLA) was performed to detect protein-protein interactions using fluorescence microscopy as previously described [61, 62]. Briefly, 293T cells were cultured in 10-chamber microscopic slides, and then co-transfected with *rev* and Keap1 or *rev* and Nrf2. 18 h later, the cells were fixed with 4% paraformaldehyde for 15min, and blocked with DuoLink blocking buffer for 1 h at 37°C. Cells were then incubated with rabbit anti-HA and mouse anti-Flag monoclonal antibodies diluted with the specific DuoLink antibody diluents for 2 h, washed for 2 minutes in 1×wash buffer A, and further incubated for 1 h at 37°C with specific PLA probes under hybridization conditions. A ligase was then added to the cells for 30 min at 37°C, to form a concatemeric product extending from the oligonucleotide arm of the PLA probe. The PLA dot was visualized as distinctly fluorescent in the Texas red channel.

### Confocal microscopy

293T cells were plated in glass coverslips on 35-mm-diameter plates. Cells were co-transfected with Nrf2 and Keap1 expression plasmids either with or without *rev*. 24 h after transfection, cells were fixed with 4% paraformaldehyde (Beyotime) for 30 min, and rinsed three times with cold phosphate-buffered saline (PBS). The fixed cells were then permeabilized for 10 min with 0.1% Triton X-100 and blocked in 5% fat-free milk in PBS for 1 h, then incubated with anti-HA, Myc or Flag tags for 1 h. Cells were washed three times with cold PBS. Then anti-mouse primary antibody, conjugated with fluorescein isothiocyanate (FITC) or Alexa Fluor 568 or Alexa Fluor 647, was added to the coverslips and the whole was incubated for 1h. Nuclei were labeled with 4′, 6-diamidino-2-phenylindole (DAPI) (Beyotime, P0131). All antibodies for immunostaining here were used at 1:500 dilution. The images were captured using a confocal microscope (LSM 880; Zeiss, Germany).

### CRISPR-Cas9 and RNAi

The Nrf2 knockout cell line was generated using CRISPR-Cas9 technology. The CRISPR-Cas9 lentiviral vector lentiCRISPR v2-cas9-GFP was purchased from Addgene (USA). The gRNA sequence used for targeting Nrf2 is GCGACGGAAAGAGTATGAGC. CRISPR/Cas9-mediated mutagenesis of Nrf2 was conducted following a previously described protocol [63]. 24 h after transfection, limiting dilution was used to allocate a single GFP-positive cell per well in 96-well plates using flow sorting. The recovered KO clones were validated using DNA sequencing and western blotting.

The Keap1 target-specific siRNAs were designed according to the Keap1 sequence and synthesized by Sigma. The nonspecific siRNA was used as a negative control to confirm the specificity of the inhibition. The 293T cells were transfected with a pool of 3 Nrf2-targeting siRNAs (TGACAGAAATTGACAACAA; AGACAAGAACAACTCCAAA; ACAGTGTCTTAACATTCAA) or control siRNA (CAAACAGAAUGGUCCUAAA) (50mM) using Lipofectamine™ RNAiMAX transfection reagent (Invitrogen, 13778100) according to the manufacturer’s instructions. The efficiency of silencing of the target gene was evaluated using western blotting.

### Protein expression and purification

cDNA encoding full-length equine Nrf2 (Genbank No. XM_008536608.1) and Keap1 (Genbank No. XM_023645423.1) were PCR-amplified from equine macrophage cells and subcloned into a pET vector containing C-terminal His6, and a PEGX6P vector with C-terminal GST tag separately. cDNA encoding EIAV *rev* was subcloned into a modified PET29a with a C-terminal MBP. For protein expression, BL21 (DE3) competent or Rosetta (DE3) competent *Escherichia coli* cells were used. Protein expression in *E. coli* cells was induced with 0.3 mM isopropyl-D-thiogalactopyranoside (IPTG) overnight at 18 °C. His-tagged Nrf2 protein and MBP-tagged Rev protein were purified with nickel affinity (HisTrap; GE Healthcare), eluted with imidazole, and desalted into binding buffer. GST-tagged Keap1 was purified on glutathione-Sepharose 4B (GE Healthcare), eluted with 10 mM reduced glutathione in 50 mM Tris, pH 8.0, and 150 mM NaCl. Protein desalinations were further performed using Akta Avant (GE Healthcare). Purity of the proteins was monitored at all stages of the purification process, and proteins were visualized using Coomassie blue staining.

### Bio-layer interferometry assay (BLI)

Binding assays were performed using an Octet Red instrument (ForteBio) based on biolayer interferometry. Streptavidin-coated biosensors were loaded with biotinylated-Keap1 in PBST (phosphate-buffered saline with 0.1 mg/ml BSA, 0.002% Tween 20, pH 7.2) at a concentration of 65μg/ml. The loaded biosensors were washed in the same buffer and then incubated together with various concentrations of Rev or Nrf2 in PBST for the indicated times to allow association. To achieve binding competition affinities, we engineered the biotinylated-Keap1 onto the streptavidin-coated biosensor first for 600 s, washed it in buffer for 180 s, and incubated it with the optimal concentrations (calculated from the affinities of Rev and Keap1) of Rev or buffer in PBST for 60 s to allow association. It was then incubated together with Nrf2. Sensors incubated together with buffer instead of Rev formed the control. The reverse competitive binding experiment was conducted in the same format. Assays were performed at room temperature in black 96-well plates (E&K Scientific). Kinetic parameters (kon and koff) and affinities (KD) were calculated using the Octet software v.6.1.

### Production of VSV-G pseudotyped retroviruses

Briefly, for rescue of the EIAV pseudovirions, 293T cells were seeded in 100-mm dishes and transfected with pONY8.1-Luc, pEIAV-GagPol and vesicular stomatitis virus glycoprotein (VSVG) expression plasmids using the PEI transfection reagent (Polysciences) as described previously. The culture supernatants were collected at 48 post-transfection and were then centrifuged at 20,000 rpm at 4 °C for 2 h. Viral pellets were resuspended in lysis buffer and subjected to western blotting analysis.

### Cell fraction and quantitative real-time PCR

293 T cells were pre-treated with Keap1 target-specific siRNA or control siRNA and then infected with EIAV pseudotyped virus. Cells were collected at 12 h and 24 h post infection. To quantify the mRNA in siKeap1 and siCon cells, total RNA from the cells was extracted using a Bio-fast simply RNA extraction kit (catalog # BSC60S1, Bioer). Nuclear and cytoplasmic RNA fractions were isolated and purified using the PARIS™ Kit (catalog # AM1921, Thermo Scientific) according to the manufacturer’s instructions. An equal volume of RNA was used for cDNA synthesis using the High-Capacity cDNA reverse transcription kit (catalog # R223-01, Vazyme Biotech). Real-time PCR was then performed using the SYBR green PCR mixture with specific primers targeting *gag*. A relative quantification method was used, with Lamin and tubulin protein as the nuclear and cytoplasmic protein controls, respectively. The following primers were used: Gag sense, 5’-GGGATTATTTGGTAAAGGG-3’, Gag anti-sense, 5’-GATTCTGCCATGCTGTTCT-3’.

### Statistics

All the graphs presented in this study were created in GraphPad Prism version 6.0 (La Jolla, CA, USA). The statistical values were calculated using one-way ANOVA or two-tailed Student’s *t*-tests. Each data bar represents the mean value ± SD (standard deviation) of at least three independent experiments in all cases. Asterisks indicate the statistical significance: ns, no significance; *, P < 0.05; **, P < 0.01.

## ACKNOWLEDGMENTS

We thank Peng Zhang for assistance on animal work. We thank the Core Facility of the Harbin Veterinary Research Institute, the Chinese Academy of Agricultural Sciences for providing the technology platform. This study was supported by grants from the National Natural Science Foundation of China (31672533 to Y L, 31302066 to X-F W). We thank OE Biotech Co., Ltd. (Shanghai,China) to help us in sequencing analysis.

## Figure legends

**S1.**
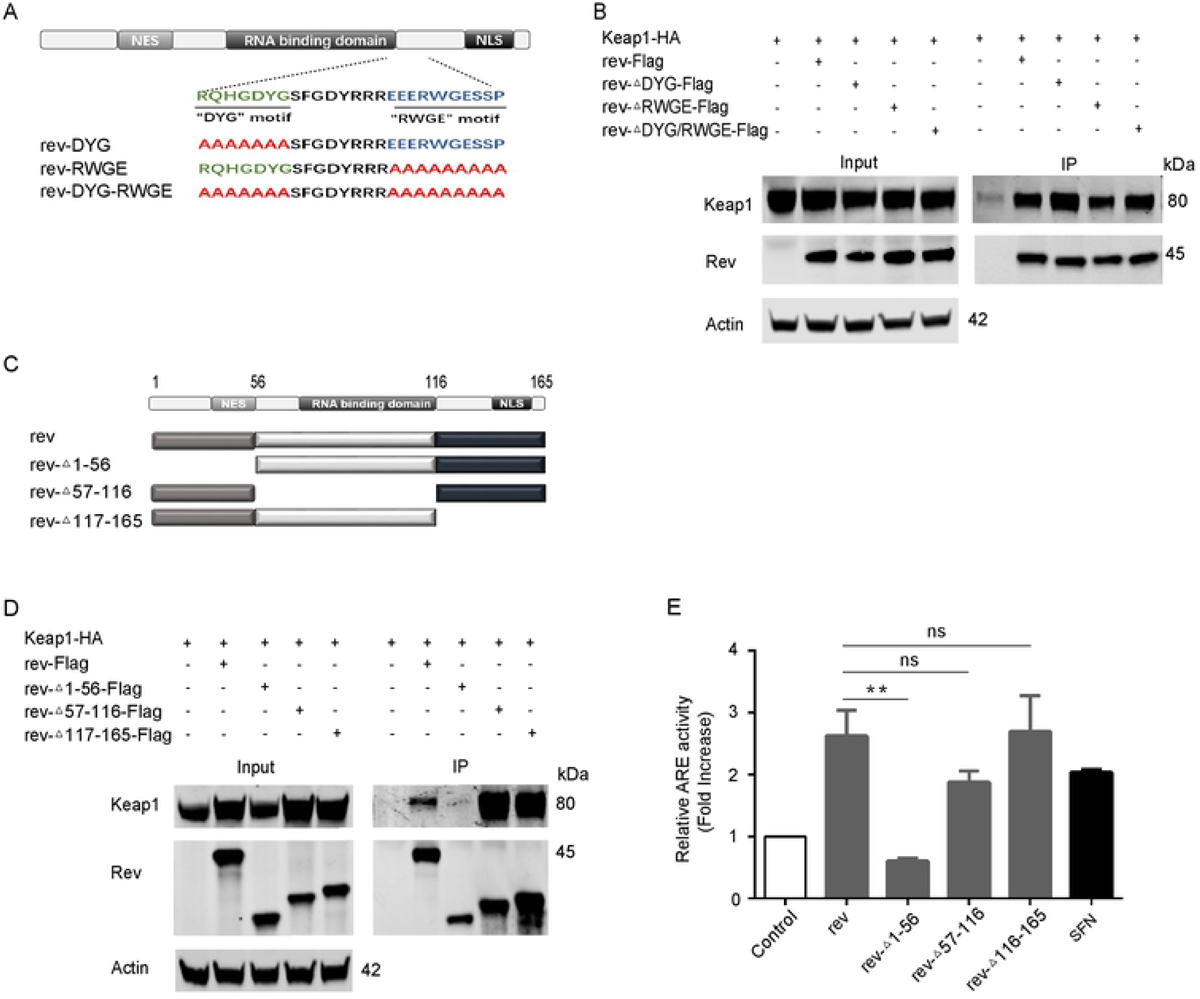
The N-terminal of EIAV-*rev* is necessary for Keap1-Rev interaction and Rev-triggered Nrf2 activation. **(A)** Schematic diagrams showing the mutations of *rev* used in B. **(B)** Western blot was performed to evaluate the interaction between Keap1 and *rev* mutants respectably. 293T cells were transfected with the indicated VR1012-based *rev* and its mutants expression plasmids with Keap1 and then analyzed by Co-IP with anti-Flag antibody. **(C)** Schematic diagram of WT EIAV-*rev* and its truncations used in D and E. **(D)** The assay protocol is as (B) but cells were transfected with *rev*-Δ1-56, *rev*-Δ57-116 or *rev*-Δ117-165 mutants. **(E)** ARE gene reporter was performed to evaluate the Nrf2/Keap1 axis activation triggered by *rev* mutants.

**S2.**
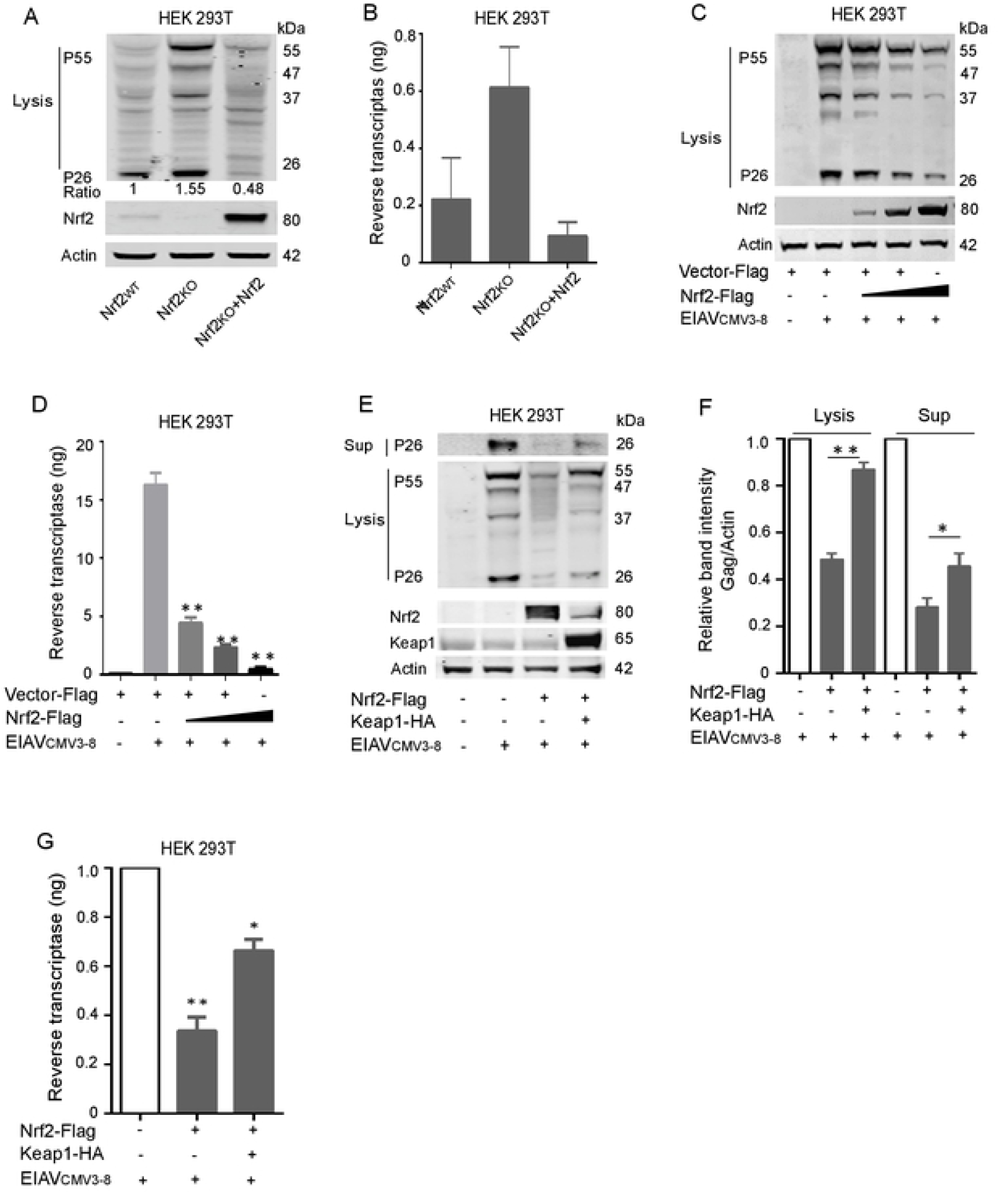
Nrf2 inhibits EIAV replication in 293T cells. **(A-B)** Viral replication was determined in 293T and Nrf2_ko_ 293T cells using western blotting (A) and reverse transcriptase activity assays (B). **(C-D)** Viral replication was evaluated in 293T cells co-transfected with EIAV_CMV3-8_ and different amounts of plasmids expressing Nrf2 respectively. Viral protein expression in virions (C) and supernatant (D) was calculated as described as A and B. **(E-G)** 293T cells were transfected with EIAV_CMV3-8_, EIAV_CMV3-8_ plus Nrf2 with or without Keap1 plasmids. Cell lysates and supernatants were analyzed as in (C-D).

## References

1. Craigo JK, Barnes S, Cook SJ, Issel CJ, Montelaro RC. Divergence, not diversity of an attenuated equine lentivirus vaccine strain correlates with protection from disease. Vaccine. 2010;28(51):8095–104. doi: 10.1016/j.vaccine.2010.10.003. PubMed PMID: 20955830; PubMed Central PMCID: PMCPMC2997116.

2. Haas L. [Equine infectious anemia--a review]. Berl Munch Tierarztl Wochenschr. 2014;127(7-8):297-300. PubMed PMID: 25080822.

3. Zimmerli U, Thur B. [Equine Infectious Anaemia - a review from an official veterinary perspective]. Schweiz Arch Tierheilkd. 2019;161(11):725-38. Epub 2019/11/07. doi: 10.17236/sat00232. PubMed PMID: 31685446.

4. Cursino AE, Vilela APP, Franco-Luiz APM, de Oliveira JG, Nogueira MF, Junior JPA, et al. Equine infectious anemia virus in naturally infected horses from the Brazilian Pantanal. Arch Virol. 2018;163(9):2385-94. Epub 2018/05/13. doi: 10.1007/s00705-018-3877-8. PubMed PMID: 29752558.

5. Wang XF, Lin YZ, Li Q, Liu Q, Zhao WW, D. C, et al. Genetic Evolution during the development of an attenuated EIAV vaccine. Retrovirology. 2016;13:9. Epub 2016/02/05. doi: 10.1186/s12977-016-0240-6. PubMed PMID: 26842878; PubMed Central PMCID: PMCPMC4738788.

6. Lin Y, Wang XF, Wang Y, Du C, Ren H, Liu C, et al. Env diversity-dependent protection of the attenuated equine infectious anaemia virus vaccine. Emerg Microbes Infect. 2020;9(1):1309-20. Epub 2020/06/12. doi: 10.1080/22221751.2020.1773323. PubMed PMID: 32525460; PubMed Central PMCID: PMCPMC7473056.

7. Liu C, Wang XF, Wang Y, Chen J, Zhong Z, Lin Y, et al. Characterization of EIAV env Quasispecies during Long-Term Passage In Vitro: Gradual Loss of Pathogenicity. Viruses. 2019;11(4). Epub 2019/04/27. doi: 10.3390/v11040380. PubMed PMID: 31022927; PubMed Central PMCID: PMCPMC6520696.

8. Whitney JB, Ruprecht RM. Live attenuated HIV vaccines: pitfalls and prospects. Curr Opin Infect Dis. 2004;17(1):17-26. PubMed PMID: 15090885.

9. Craigo JK, Montelaro RC. Equine Infectious Anemia Virus Infection and Immunity: Lessons for Aids Vaccine Development. Future Virol. 2011;6(2):139–42. doi: 10.2217/fvl.10.85. PubMed PMID: 21643555; PubMed Central PMCID: PMCPMC3107501.

10. Ji S, Na L, Ren H, Wang Y, Wang X. Equine Myxovirus Resistance Protein 2 Restricts Lentiviral Replication by Blocking Nuclear Uptake of Capsid Protein. J Virol. 2018;92(18). Epub 2018/05/11. doi: 10.1128/JVI.00499-18. PubMed PMID: 29743377; PubMed Central PMCID: PMCPMC6146692.

11. Lin YZ, Sun LK, Zhu DT, Hu Z, Wang XF, D. C, et al. Equine schlafen 11 restricts the production of equine infectious anemia virus via a codon usage-dependent mechanism. Virology. 2016;495:112-21. Epub 2016/05/21. doi: 10.1016/j.virol.2016.04.024. PubMed PMID: 27200480.

12. Yin X, Hu Z, Gu Q, Wu X, Zheng YH, Wei P, et al. Equine tetherin blocks retrovirus release and its activity is antagonized by equine infectious anemia virus envelope protein. J Virol. 2014;88(2):1259-70. Epub 2013/11/15. doi: 10.1128/JVI.03148-13. PubMed PMID: 24227834; PubMed Central PMCID: PMCPMC3911658.

13. Ren H, Yin X, Su C, Guo M, Wang XF, Na L, et al. Equine lentivirus counteracts SAMHD1 restriction by Rev-mediated degradation of SAMHD1 via the BECN1-dependent lysosomal pathway. Autophagy. 2020:1-18. Epub 2020/11/12. doi: 10.1080/15548627.2020.1846301. PubMed PMID: 33172327.

14. Fraternale A, Zara C, De Angelis M, Nencioni L, Palamara AT, Retini M, et al. Intracellular Redox-Modulated Pathways as Targets for Effective Approaches in the Treatment of Viral Infection. Int J Mol Sci. 2021;22(7). Epub 2021/04/04. doi: 10.3390/ijms22073603. PubMed PMID: 33808471; PubMed Central PMCID: PMCPMC8036776.

15. Antonucci S, Di Lisa F, Kaludercic N. Mitochondrial reactive oxygen species in physiology and disease. Cell Calcium. 2021;94:102344. Epub 2021/02/09. doi: 10.1016/j.ceca.2020.102344. PubMed PMID: 33556741.

16. Zevini A, Ferrari M, Olagnier D, Hiscott J. Dengue virus infection and Nrf2 regulation of oxidative stress. Curr Opin Virol. 2020;43:35-40. Epub 2020/08/24. doi: 10.1016/j.coviro.2020.07.015. PubMed PMID: 32829129.

17. Sharma N, Muthamilarasan M, Dulani P, Prasad M. Genomic dissection of ROS detoxifying enzyme encoding genes for their role in antioxidative defense mechanism against Tomato leaf curl New Delhi virus infection in tomato. Genomics. 2021. Epub 2021/02/02. doi: 10.1016/j.ygeno.2021.01.022. PubMed PMID: 33524498.

18. Pan YG, Huang MT, Sekar P, Huang DY, Lin WW, Hsieh SL. Decoy Receptor 3 Inhibits Monosodium Urate-Induced NLRP3 Inflammasome Activation via Reduction of Reactive Oxygen Species Production and Lysosomal Rupture. Front Immunol. 2021;12:638676. Epub 2021/03/23. doi: 10.3389/fimmu.2021.638676. PubMed PMID: 33746978; PubMed Central PMCID: PMCPMC7966727.

19. Ren CZ, Hu WY, Zhang JW, Wei YY, Yu ML, Hu TJ. Establishment of inflammatory model induced by Pseudorabies virus infection in mice. J Vet Sci. 2021;22(2):e20. Epub 2021/03/29. doi: 10.4142/jvs.2021.22.e20. PubMed PMID: 33774936; PubMed Central PMCID: PMCPMC8007442.

20. Kansanen E, Kuosmanen SM, Leinonen H, Levonen AL. The Keap1-Nrf2 pathway: Mechanisms of activation and dysregulation in cancer. Redox Biol. 2013;1:45-9. Epub 2013/09/12. doi: 10.1016/j.redox.2012.10.001. PubMed PMID: 24024136; PubMed Central PMCID: PMCPMC3757665.

21. Shaw P, Chattopadhyay A. Nrf2-ARE signaling in cellular protection: Mechanism of action and the regulatory mechanisms. J Cell Physiol. 2020;235(4):3119-30. Epub 2019/09/25. doi: 10.1002/jcp.29219. PubMed PMID: 31549397.

22. Lee C. Therapeutic Modulation of Virus-Induced Oxidative Stress via the Nrf2-Dependent Antioxidative Pathway. Oxid Med Cell Longev. 2018;2018:6208067. Epub 2018/12/06. doi: 10.1155/2018/6208067. PubMed PMID: 30515256; PubMed Central PMCID: PMCPMC6234444.

23. Mohan S, Gupta D. Crosstalk of toll-like receptors signaling and Nrf2 pathway for regulation of inflammation. Biomed Pharmacother. 2018;108:1866-78. Epub 2018/10/31. doi: 10.1016/j.biopha.2018.10.019. PubMed PMID: 30372892.

24. Mu S, Yang W, Huang G. Antioxidant activities and mechanisms of polysaccharides. Chem Biol Drug Des. 2020. Epub 2020/09/19. doi: 10.1111/cbdd.13798. PubMed PMID: 32946177.

25. Olagnier D, Brandtoft AM, Gunderstofte C, Villadsen NL, Krapp C, Thielke AL, et al. Nrf2 negatively regulates STING indicating a link between antiviral sensing and metabolic reprogramming. Nat Commun. 2018;9(1):3506. Epub 2018/08/31. doi: 10.1038/s41467-018-05861-7. PubMed PMID: 30158636; PubMed Central PMCID: PMCPMC6115435.

26. Okawa H, Motohashi H, Kobayashi A, Aburatani H, Kensler TW, Yamamoto M. Hepatocyte-specific deletion of the keap1 gene activates Nrf2 and confers potent resistance against acute drug toxicity. Biochem Biophys Res Commun. 2006;339(1):79-88. Epub 2005/11/19. doi: 10.1016/j.bbrc.2005.10.185. PubMed PMID: 16293230.

27. Deshmukh P, Unni S, Krishnappa G, Padmanabhan B. The Keap1-Nrf2 pathway: promising therapeutic target to counteract ROS-mediated damage in cancers and neurodegenerative diseases. Biophys Rev. 2017;9(1):41-56. Epub 2017/05/17. doi: 10.1007/s12551-016-0244-4. PubMed PMID: 28510041; PubMed Central PMCID: PMCPMC5425799.

28. Lee JM, Johnson JA. An important role of Nrf2-ARE pathway in the cellular defense mechanism. J Biochem Mol Biol. 2004;37(2):139-43. Epub 2004/10/08. doi: 10.5483/bmbrep.2004.37.2.139. PubMed PMID: 15469687.

29. Shao J, Huang J, Guo Y, Li L, Liu X, Chen X, et al. Up-regulation of nuclear factor E2-related factor 2 (Nrf2) represses the replication of SVCV. Fish Shellfish Immunol. 2016;58:474-82. Epub 2016/10/23. doi: 10.1016/j.fsi.2016.09.012. PubMed PMID: 27693327.

30. Page A, Volchkova VA, Reid SP, Mateo M, Bagnaud-Baule A, Nemirov K, et al. Marburgvirus hijacks nrf2-dependent pathway by targeting nrf2-negative regulator keap1. Cell Rep. 2014;6(6):1026-36. Epub 2014/03/19. doi: 10.1016/j.celrep.2014.02.027. PubMed PMID: 24630992.

31. Kosmider B, Messier EM, Janssen WJ, Nahreini P, Wang J, Hartshorn KL, et al. Nrf2 protects human alveolar epithelial cells against injury induced by influenza A virus. Respir Res. 2012;13:43. Epub 2012/06/08. doi: 10.1186/1465-9921-13-43. PubMed PMID: 22672594; PubMed Central PMCID: PMCPMC3520784.

32. Saeedi-Boroujeni A, Mahmoudian-Sani MR. Anti-inflammatory potential of Quercetin in COVID-19 treatment. J Inflamm (Lond). 2021;18(1):3. Epub 2021/01/30. doi: 10.1186/s12950-021-00268-6. PubMed PMID: 33509217; PubMed Central PMCID: PMCPMC7840793.

33. Lin YZ, Cao XZ, Li L, Li L, Jiang CG, Wang XF, et al. The pathogenic and vaccine strains of equine infectious anemia virus differentially induce cytokine and chemokine expression and apoptosis in macrophages. Virus Res. 2011;160(1-2):274-82. Epub 2011/07/26. doi: 10.1016/j.virusres.2011.06.028. PubMed PMID: 21782860.

34. Du C, Duan Y, Wang XF, Lin Y, Na L, Wang X, et al. Attenuation of Equine Lentivirus Alters Mitochondrial Protein Expression Profile from Inflammation to Apoptosis. J Virol. 2019;93(21). Epub 2019/08/09. doi: 10.1128/JVI.00653-19. PubMed PMID: 31391270; PubMed Central PMCID: PMCPMC6803291.

35. Kobayashi A, Kang MI, Watai Y, Tong KI, Shibata T, Uchida K, et al. Oxidative and electrophilic stresses activate Nrf2 through inhibition of ubiquitination activity of Keap1. Mol Cell Biol. 2006;26(1):221-9. Epub 2005/12/16. doi: 10.1128/MCB.26.1.221-229.2006. PubMed PMID: 16354693; PubMed Central PMCID: PMCPMC1317630.

36. Watts NR, Eren E, Zhuang X, Wang YX, Steven AC, Wingfield PT. A new HIV-1 Rev structure optimizes interaction with target RNA (RRE) for nuclear export. J Struct Biol. 2018;203(2):102-8. Epub 2018/04/02. doi: 10.1016/j.jsb.2018.03.011. PubMed PMID: 29605570; PubMed Central PMCID: PMCPMC6019186.

37. Bai Y, Tambe A, Zhou K, Doudna JA. RNA-guided assembly of Rev-RRE nuclear export complexes. Elife. 2014;3:e03656. Epub 2014/08/29. doi: 10.7554/eLife.03656. PubMed PMID: 25163983; PubMed Central PMCID: PMCPMC4142337.

38. Ramezani A, Nahad MP, Faghihloo E. The role of Nrf2 transcription factor in viral infection. J Cell Biochem. 2018;119(8):6366-82. Epub 2018/05/09. doi: 10.1002/jcb.26897. PubMed PMID: 29737559.

39. Bender D, Hildt E. Effect of Hepatitis Viruses on the Nrf2/Keap1-Signaling Pathway and Its Impact on Viral Replication and Pathogenesis. Int J Mol Sci. 2019;20(18). Epub 2019/09/25. doi: 10.3390/ijms20184659. PubMed PMID: 31546975; PubMed Central PMCID: PMCPMC6769940.

40. Fan X, Murray SC, Staitieh BS, Spearman P, Guidot DM. HIV Impairs Alveolar Macrophage Function via MicroRNA-144-Induced Suppression of Nrf2. Am J Med Sci. 2021;361(1):90-7. Epub 2020/08/11. doi: 10.1016/j.amjms.2020.07.026. PubMed PMID: 32773107; PubMed Central PMCID: PMCPMC7854972.

41. Gjyshi O, Bottero V, Veettil MV, Dutta S, Singh VV, Chikoti L, et al. Kaposi’s sarcoma-associated herpesvirus induces Nrf2 during de novo infection of endothelial cells to create a microenvironment conducive to infection. PLoS Pathog. 2014;10(10):e1004460. Epub 2014/10/24. doi: 10.1371/journal.ppat.1004460. PubMed PMID: 25340789; PubMed Central PMCID: PMCPMC4207826.

42. Gjyshi O, Roy A, Dutta S, Veettil MV, Dutta D, Chandran B. Activated Nrf2 Interacts with Kaposi’s Sarcoma-Associated Herpesvirus Latency Protein LANA-1 and Host Protein KAP1 To Mediate Global Lytic Gene Repression. J Virol. 2015;89(15):7874-92. Epub 2015/05/23. doi: 10.1128/JVI.00895-15. PubMed PMID: 25995248; PubMed Central PMCID: PMCPMC4505678.

43. Gjyshi O, Flaherty S, Veettil MV, Johnson KE, Chandran B, Bottero V. Kaposi’s sarcoma-associated herpesvirus induces Nrf2 activation in latently infected endothelial cells through SQSTM1 phosphorylation and interaction with polyubiquitinated Keap1. J Virol. 2015;89(4):2268-86. Epub 2014/12/17. doi: 10.1128/JVI.02742-14. PubMed PMID: 25505069; PubMed Central PMCID: PMCPMC4338888.

44. Komaravelli N, Tian B, Ivanciuc T, Mautemps N, Brasier AR, Garofalo RP, et al. Respiratory syncytial virus infection down-regulates antioxidant enzyme expression by triggering deacetylation-proteasomal degradation of Nrf2. Free Radic Biol Med. 2015;88(Pt B):391-403. Epub 2015/06/16. doi: 10.1016/j.freeradbiomed.2015.05.043. PubMed PMID: 26073125; PubMed Central PMCID: PMCPMC4628892.

45. Komaravelli N, Ansar M, Garofalo RP, Casola A. Respiratory syncytial virus induces NRF2 degradation through a promyelocytic leukemia protein - ring finger protein 4 dependent pathway. Free Radic Biol Med. 2017;113:494-504. Epub 2017/11/07. doi: 10.1016/j.freeradbiomed.2017.10.380. PubMed PMID: 29107745; PubMed Central PMCID: PMCPMC5699968.

46. Ferrari M, Zevini A, Palermo E, Muscolini M, Alexandridi M, Etna MP, et al. Dengue Virus Targets Nrf2 for NS2B3-Mediated Degradation Leading to Enhanced Oxidative Stress and Viral Replication. J Virol. 2020;94(24). Epub 2020/10/02. doi: 10.1128/jvi.01551-20. PubMed PMID: 32999020; PubMed Central PMCID: PMCPMC7925186.

47. Edwards MR, Johnson B, Mire CE, Xu W, Shabman RS, Speller LN, et al. The Marburg virus VP24 protein interacts with Keap1 to activate the cytoprotective antioxidant response pathway. Cell Rep. 2014;6(6):1017-25. Epub 2014/03/19. doi: 10.1016/j.celrep.2014.01.043. PubMed PMID: 24630991; PubMed Central PMCID: PMCPMC3985291.

48. Mastrantonio R, Cervelli M, Pietropaoli S, Mariottini P, Colasanti M, Persichini T. HIV-Tat Induces the Nrf2/ARE Pathway through NMDA Receptor-Elicited Spermine Oxidase Activation in Human Neuroblastoma Cells. PLoS One. 2016;11(2):e0149802. Epub 2016/02/20. doi: 10.1371/journal.pone.0149802. PubMed PMID: 26895301; PubMed Central PMCID: PMCPMC4760771.

49. Silva-Islas CA, Maldonado PD. Canonical and non-canonical mechanisms of Nrf2 activation. Pharmacol Res. 2018;134:92-9. Epub 2018/06/19. doi: 10.1016/j.phrs.2018.06.013. PubMed PMID: 29913224.

50. Dodson M, Zhang DD. Non-canonical activation of NRF2: New insights and its relevance to disease. Curr Pathobiol Rep. 2017;5(2):171-6. Epub 2017/10/31. doi: 10.1007/s40139-017-0131-0. PubMed PMID: 29082113; PubMed Central PMCID: PMCPMC5654572.

51. Zhang DD, Hannink M. Distinct cysteine residues in Keap1 are required for Keap1-dependent ubiquitination of Nrf2 and for stabilization of Nrf2 by chemopreventive agents and oxidative stress. Mol Cell Biol. 2003;23(22):8137-51. Epub 2003/10/31. doi: 10.1128/mcb.23.22.8137-8151.2003. PubMed PMID: 14585973; PubMed Central PMCID: PMCPMC262403.

52. Saito Y, Yako T, Otsu W, Nakamura S, Inoue Y, Muramatsu A, et al. A triterpenoid Nrf2 activator, RS9, promotes LC3-associated phagocytosis of photoreceptor outer segments in a p62-independent manner. Free Radic Biol Med. 2020;152:235-47. Epub 2020/03/29. doi: 10.1016/j.freeradbiomed.2020.03.012. PubMed PMID: 32217192.

53. Hartwick Bjorkman S, Oliveira Pereira R. The Interplay Between Mitochondrial Reactive Oxygen Species, Endoplasmic Reticulum Stress, and Nrf2 Signaling in Cardiometabolic Health. Antioxid Redox Signal. 2021. Epub 2021/02/19. doi: 10.1089/ars.2020.8220. PubMed PMID: 33599550.

54. Tong KI, Kobayashi A, Katsuoka F, Yamamoto M. Two-site substrate recognition model for the Keap1-Nrf2 system: a hinge and latch mechanism. Biol Chem. 2006;387(10-11):1311-20. Epub 2006/11/04. doi: 10.1515/BC.2006.164. PubMed PMID: 17081101.

55. Tong KI, Katoh Y, Kusunoki H, Itoh K, Tanaka T, Yamamoto M. Keap1 recruits Neh2 through binding to ETGE and DLG motifs: characterization of the two-site molecular recognition model. Mol Cell Biol. 2006;26(8):2887-900. Epub 2006/04/04. doi: 10.1128/MCB.26.8.2887-2900.2006. PubMed PMID: 16581765; PubMed Central PMCID: PMCPMC1446969.

56. Hassan SM, Jawad MJ, Ahjel SW, Singh RB, Singh J, Awad SM, et al. The Nrf2 Activator (DMF) and Covid-19: Is there a Possible Role? Med Arch. 2020;74(2):134-8. Epub 2020/06/25. doi: 10.5455/medarh.2020.74.134-138. PubMed PMID: 32577056; PubMed Central PMCID: PMCPMC7296400.

57. McCord JM, Hybertson BM, Cota-Gomez A, Gao B. Nrf2 Activator PB125(R) as a Potential Therapeutic Agent Against COVID-19. bioRxiv. 2020. Epub 2020/06/09. doi: 10.1101/2020.05.16.099788. PubMed PMID: 32511372; PubMed Central PMCID: PMCPMC7263501.

58. Bolger AM, Lohse M, Usadel B. Trimmomatic: a flexible trimmer for Illumina sequence data. Bioinformatics. 2014;30(15):2114-20. Epub 2014/04/04. doi: 10.1093/bioinformatics/btu170. PubMed PMID: 24695404; PubMed Central PMCID: PMCPMC4103590.

59. Anders S, Pyl PT, Huber W. HTSeq--a Python framework to work with high-throughput sequencing data. Bioinformatics. 2015;31(2):166-9. Epub 2014/09/28. doi: 10.1093/bioinformatics/btu638. PubMed PMID: 25260700; PubMed Central PMCID: PMCPMC4287950.

60. Xu Q, Zhao W, Chen Y, Tong Y, Rong G, Huang Z, et al. Transcriptome profiling of the goose (Anser cygnoides) ovaries identify laying and broodiness phenotypes. PLoS One. 2013;8(2):e55496. Epub 2013/02/14. doi: 10.1371/journal.pone.0055496. PubMed PMID: 23405160; PubMed Central PMCID: PMCPMC3566205.

61. Gullberg M, Gustafsdottir SM, Schallmeiner E, Jarvius J, Bjarnegard M, Betsholtz C, et al. Cytokine detection by antibody-based proximity ligation. Proc Natl Acad Sci U S A. 2004;101(22):8420-4. Epub 2004/05/25. doi: 10.1073/pnas.0400552101. PubMed PMID: 15155907; PubMed Central PMCID: PMCPMC420409.

62. Andersen SS, Hvid M, Pedersen FS, Deleuran B. Proximity ligation assay combined with flow cytometry is a powerful tool for the detection of cytokine receptor dimerization. Cytokine. 2013;64(1):54-7. Epub 2013/06/04. doi: 10.1016/j.cyto.2013.04.026. PubMed PMID: 23726671.

63. Yasuda K, Nishikawa M, Okamoto K, Horibe K, Mano H, Yamaguchi M, et al. Elucidation of metabolic pathways of 25-hydroxyvitamin D3 mediated by Cyp24a1 and Cyp3a using Cyp24a1 knockout rats generated by CRISPR/Cas9 system. J Biol Chem. 2021:100668. Epub 2021/04/19. doi: 10.1016/j.jbc.2021.100668. PubMed PMID: 33865853.

